# Calcineurin associates with centrosomes and regulates cilia length maintenance

**DOI:** 10.1101/2022.06.16.496489

**Authors:** Eirini Tsekitsidou, Jennifer T. Wang, Cassandra J. Wong, Idil Ulengin-Talkish, Tim Stearns, Anne-Claude Gingras, Martha S. Cyert

## Abstract

Calcineurin, or PP2B, the Ca^2+^ and calmodulin-activated phosphatase and target of immunosuppressants, has many substrates and functions that remain undiscovered. By combining rapid proximity-dependent labeling with cell cycle synchronization, we mapped calcineurin’s spatial distribution in different cell cycle stages. While calcineurin-proximal proteins did not vary significantly between interphase and mitosis, calcineurin consistently associated with multiple centrosomal/ciliary proteins. These include POC5, which binds centrin in a Ca^2+^-dependent manner and is a component of the luminal scaffold that stabilizes centrioles. We show that POC5 contains a calcineurin substrate motif (PxIxIT-type) that mediates calcineurin binding *in vivo* and *in vitro*. Using indirect immunofluorescence and expansion microscopy, we demonstrate that calcineurin co-localizes with POC5 at the centrosome, and further show that calcineurin inhibitors alter POC5 distribution within the centriole lumen. Our discovery that calcineurin directly associates with centrosomal proteins highlights a role for Ca^2+^ and calcineurin signaling at these organelles. Calcineurin inhibition promotes primary cilia elongation without affecting ciliogenesis. Thus, Ca^2+^ signaling within cilia includes previously unknown functions for calcineurin in cilia length maintenance, a process frequently disrupted in ciliopathies.

**Summary statement:** Calcineurin phosphatase participates in centrosome and cilia regulation. Calcineurin localizes to centrosomes, where it interacts with partner POC5, and its inhibition promotes cilia elongation.

## Introduction

In cells, calcium signaling is spatially controlled through colocalization of calcium ions (Ca^2+^) with their effector proteins within microdomains, allowing Ca^2+^ signals of different origins to direct distinct downstream events (Berridge et al., 2003). One such effector is calcineurin (CN, also known as protein phosphatase 2B or PP2B), the sole Ca^2+^/calmodulin (CaM) regulated serine/threonine protein phosphatase in animals. CN activates the adaptive immune response by dephosphorylating NFAT (nuclear factor of activated T-cells) transcription factors in T-cells, and CN inhibitors (CNIs) FK506/tacrolimus and cyclosporin A (CysA), are used clinically as immunosuppressants (Rusnak and Mertz, 2000). However, CN is ubiquitously expressed, and has demonstrated roles in the cardiovascular and nervous systems. Thus CNIs also cause a broad range of adverse effects, especially in the kidney, whose etiologies are largely unknown (Azzi et al., 2013; Farouk and Rein, 2020). This underscores the need to elucidate CN signaling throughout the body, and to understand its targeting to different subcellular locations.

CN is an obligate heterodimer composed of a regulatory (CNB) and catalytic (CNA) subunit, which is inactive until Ca^2+^/CaM binds to CNA and relieves autoinhibition (Li et al., 2016; Rusnak and Mertz, 2000). CN recognizes substrates via two **<u>S</u>**hort **<u>Li</u>**near peptide **<u>M</u>**otifs, or SLiMs, i.e. degenerate 3- to 10-aa sequences found within intrinsically disordered domains, that mediate low affinity, dynamic protein-protein interactions during signaling (Tompa et al., 2014). The CN-binding SLiMs, PxIxIT and LxVP, have distinct properties: LxVP motifs bind to a pocket that is accessible only in the activated enzyme and is targeted by CNIs to block substrate dephosphorylation (Grigoriu et al., 2013; Roy and Cyert, 2020). In contrast, PxIxIT motifs bind directly to CNA independently of its activation state and determine CN’s intracellular distribution via anchoring to substrates, regulators and scaffolds (Roy and Cyert, 2020).

CN-specific SLiMs have been leveraged to systematically decode CN signaling. *In silico* identification of PxIxIT and LxVP motifs revealed hundreds of putative substrates in humans (Brauer et al., 2019; Sheftic et al., 2016; Wigington et al., 2020). Experimentally, weak affinity SLiM-dependent CN interactions have been captured using proximity-dependent biotinylation coupled to mass spectrometry (PDB-MS) (Ulengin-Talkish et al., 2021; Wigington et al., 2020). Fusion of either WT or mutant CNAs (defective for PxIxIT or LxVP binding) to the promiscuous biotin ligase, BirA*, identified CN-proximal proteins. 50% of these were SLiM-dependent, i.e. showed reduced labeling with mutant CNs, and were enriched for computationally predicted CN-binding SLiMs. In humans, CN-proximal proteins map to multiple cellular compartments, which surprisingly include centrosomes, where its functions are unknown (Wigington et al., 2020).

Centrosomes serve as the cell’s microtubule organizing center (MTOC), and are membrane-less organelles formed by two centrioles associated with pericentriolar material (PCM). The centrosome nucleates microtubules both in interphase and in mitosis, when duplicated centrosomes form the poles of the mitotic spindle (Wang and Stearns, 2017). In most cells that are not actively proliferating, centrioles transform into basal bodies which direct formation of the primary cilium, a non-motile appendage that extends into the extracellular space and functions as the cell’s molecular antenna. (Avasthi and Marshall, 2012; Wang and Stearns, 2017). Primary cilia serve as specialized sites for Hedgehog and Ca^2+^ signaling, and when disrupted, lead to a group of disorders known as ciliopathies (Delling et al., 2013; Hildebrandt et al., 2011).

Here, we further investigate the association of CN with centrosomes and cilia by mapping CN-proximal proteins using miniTurbo, a fast-acting biotin ligase whose short labelling time allowed us to probe CN’s subcellular distribution across the cell cycle. (Branon et al., 2018). We find that CN-proximal proteins do not change dramatically between interphase and mitosis, but are significantly enriched at centrosomes and centrioles, including POC5 (proteome of centriole 5), which we demonstrate to bind CN via a PxIxIT motif. We show that a pool of CN colocalizes with POC5 at centrosomes, and that CNIs alter POC5 distribution within the centriole. Finally, we demonstrate that CN inhibition promotes primary cilia elongation without affecting ciliogenesis. Together, our findings establish that CN associates directly with centrosomal components and regulates cilia length, a process frequently disrupted in ciliopathies.

## Results and Discussion

### Proximity labeling maps CN’s subcellular distribution across the cell cycle

To map CN’s subcellular neighborhoods across the cell cycle, CNAα was fused to miniTurbo, a promiscuous biotin ligase that labels proteins within a ∼10nm radius in 15 minutes (Branon et al., 2018; Gingras et al., 2019). HEK293 T-REx cells expressing miniTurbo-3xFLAG alone or fused to -CNAα_WT_ or -CNAα_NIRmut_, a mutant with impaired PxIxIT-docking (Li et al., 2007), were incubated with biotin using populations that were either asynchronous, or synchronized in G1/S (released from double thymidine block) or mitosis (released from arrest with Cdk1 inhibitor RO-3306) (Fig. 1A). MiniTurbo-CNA expression did not alter cell cycle progression, and cell synchrony and miniTurbo expression levels were evaluated for each sample (Figs S1A-E).

**Fig. 1.**
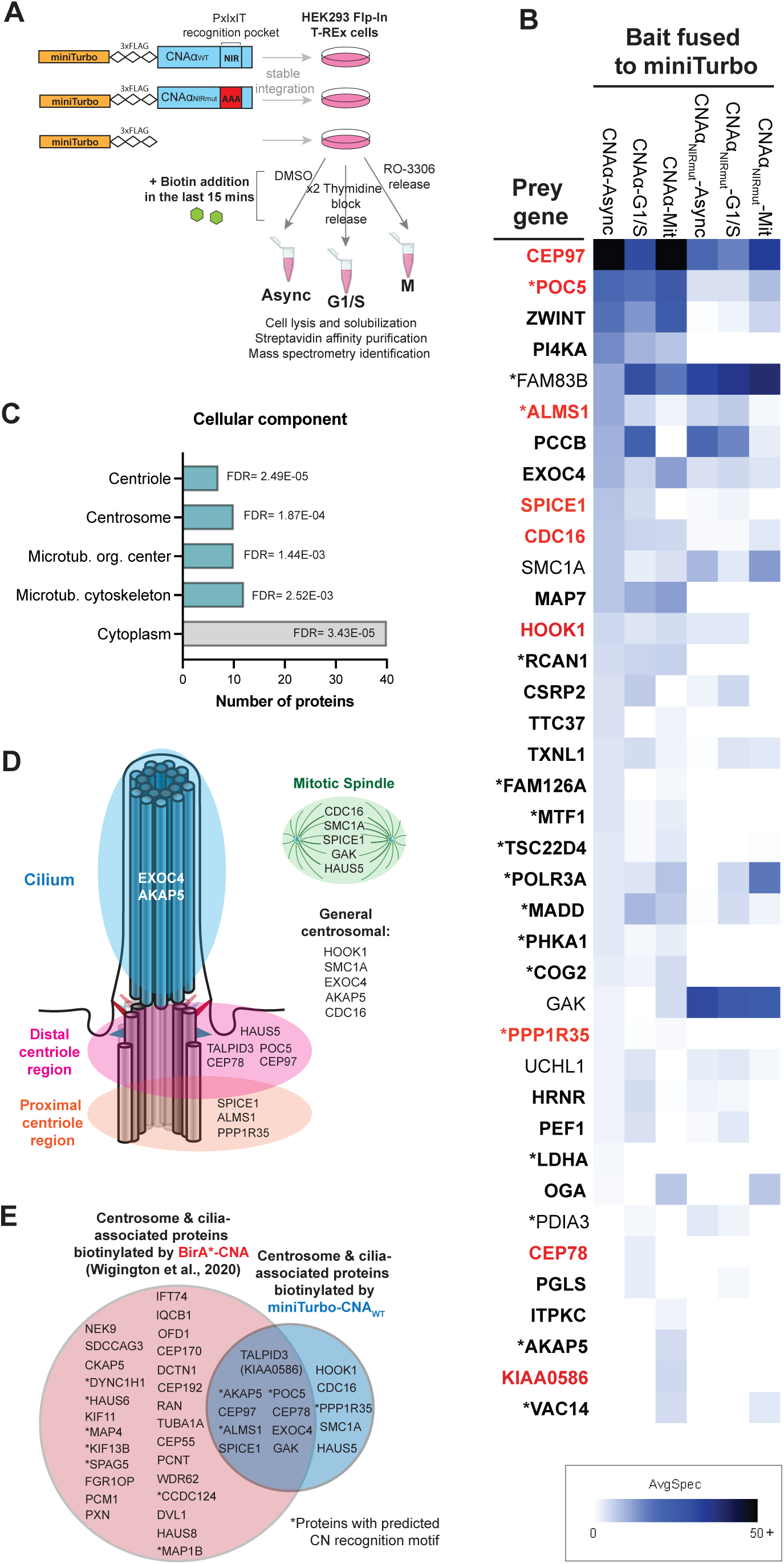
Temporal mapping of calcineurin-proximal proteins reveals centrosome association. A. Scheme for proximity-dependent biotin labeling using asynchronous or synchronized cells expressing miniTurbo-3xFLAG alone or fused to CNA_WT_ or CNA_NIRmut_. B. Average spectral counts determined via DDA for preys labelled by CNA_WT_ with ≥ 2 peptides and BFDR ≤ 0.01 in at least one condition. Bold: PxIxIT-dependent (Log_2_ CNA_WT/_CNA_NIRmut_ ≥ 0.5). Red: Preys from GO category “centrosome”. Asterisks: preys with predicted CN-dependent SLiMs. C. Select statistically enriched GO cellular component categories shown for the 41 CN-proximal proteins. FDR, false discovery rate. D. Schematic showing locations CN-proximal proteins that localize to centrosomes, cilia or mitotic spindles, details in Table S2. E. Overlap of centrosome and cilia-associated CN-proximal proteins detected by Wigington et al., 2020 (pink circle) or in this study (blue). Asterisks indicate proteins with predicted CN-binding SLiMs.

Biotinylated proteins were then identified via mass spectrometry (MS) using both data-dependent (DDA) and data-independent acquisition (DIA/mSPLIT) which yielded 41 CN-proximal proteins that were significantly biotinylated by CNAα_WT_ (≥ 2 unique peptides and bayesian false discovery rate, BFDR ≤ 0.01) in at least one condition (38 from DDA and 3 additional from mSPLIT-Figs 1B, S1F and Table S1). These proteins are enriched for protein-protein interactions (PPI enrichment p-value = 4.02e-05, Fig. S1G) and include several known CN interactors and substrates: AKAP5 (Dell’Acqua et al., 2002), PI4KA and FAM126A (Ulengin-Talkish et al., 2021), PHKA1 (Ingebritsen and Cohen, 1983) and RCAN1 (Mehta et al., 2009). The majority of CN-proximal proteins were PxIxIT-dependent (35/41, Log_2_ spectral counts with CNAα_WT_/CNAα_NIRmut_ ≥ 0.5 for at least one condition), and 17 of these contained a predicted CN-specific SLiM (Figs 1B, S1F and Table S1). Fewer CN-proximal proteins were identified compared to previous studies with BirA* (Wigington et al., 2020), likely due to the much shorter labeling time (15 minutes versus 18 hours). However, 15 proteins were common to both datasets (Table S1).

Notably, the most robustly detected CN-proximal proteins were common to asynchronous, G1/S and mitotic populations (23/41 preys). Proteins that were detected only in M (4/41) or G1/S (3/41) samples were generally not known to be associated with cell-cycle-specific functions and were represented by relatively few spectral counts (≤7), suggesting that they may be at the limit of detection. Together, these observations suggest that CN spatial distribution is relatively constant throughout the cell cycle and that any changes in CN signaling are dictated instead by temporal regulation of Ca^2+^ signals.

### CN is proximal to centrosomal components in a cell cycle independent manner

Interestingly, CN-proximal proteins identified here were significantly enriched in gene ontology terms associated with the microtubule cytoskeleton, centrosomes and specifically centrioles, with the reactome pathway “cilium assembly” also being enriched (FDR 0.044) (Mi et al., 2013) (Figs 1B, C, Table S1). Furthermore, by manually curating the literature we found 14 CN-proximal proteins that spatially distribute throughout the centrosome and cilium and/or localize to the mitotic spindle (Fig. 1D and Table S2). Nine of these were identified previously (Fig. 1E).

Overall, these results suggest that CN contacts centrosomal proteins, either at the centrosome or before they are incorporated into centrosomes. In addition, CN proximity to mitotic spindle proteins suggests possible undiscovered functions for CN at spindle microtubules or poles during mitosis, when centrosomal Ca^2+^ signals have been observed (Helassa et al., 2019).

### CN regulatory subunit CNB localizes to the centrosome

Our evidence of CN proximity to centrosomal/ciliary proteins spurred us to examine whether a pool of CN exists at these organelles. Initial attempts to visualize CNB (which is always found in association with CNA) via indirect immunofluorescence of hTERT-RPE1 showed diffuse cytoplasmic distribution.

However, by permeabilizing cells with digitonin treatment prior to fixation (Sydor et al., 2018), we were able to detect CNB localization at centrioles and the pericentriolar region of centrosomes (Fig. 2A). Specifically, CNB was observed either surrounding centrioles (63.9% of cells) or distributed in between mother and daughter centrioles (25.3% of cells), with only 10.8% of cells showing no centriole-associated CNB. Furthermore, this staining was specific, as centrosomal localization was eliminated by preincubating the anti-CNB antisera with purified CNB protein, but not with bovine serum albumin (Figs S2A, B).

**Fig. 2.**
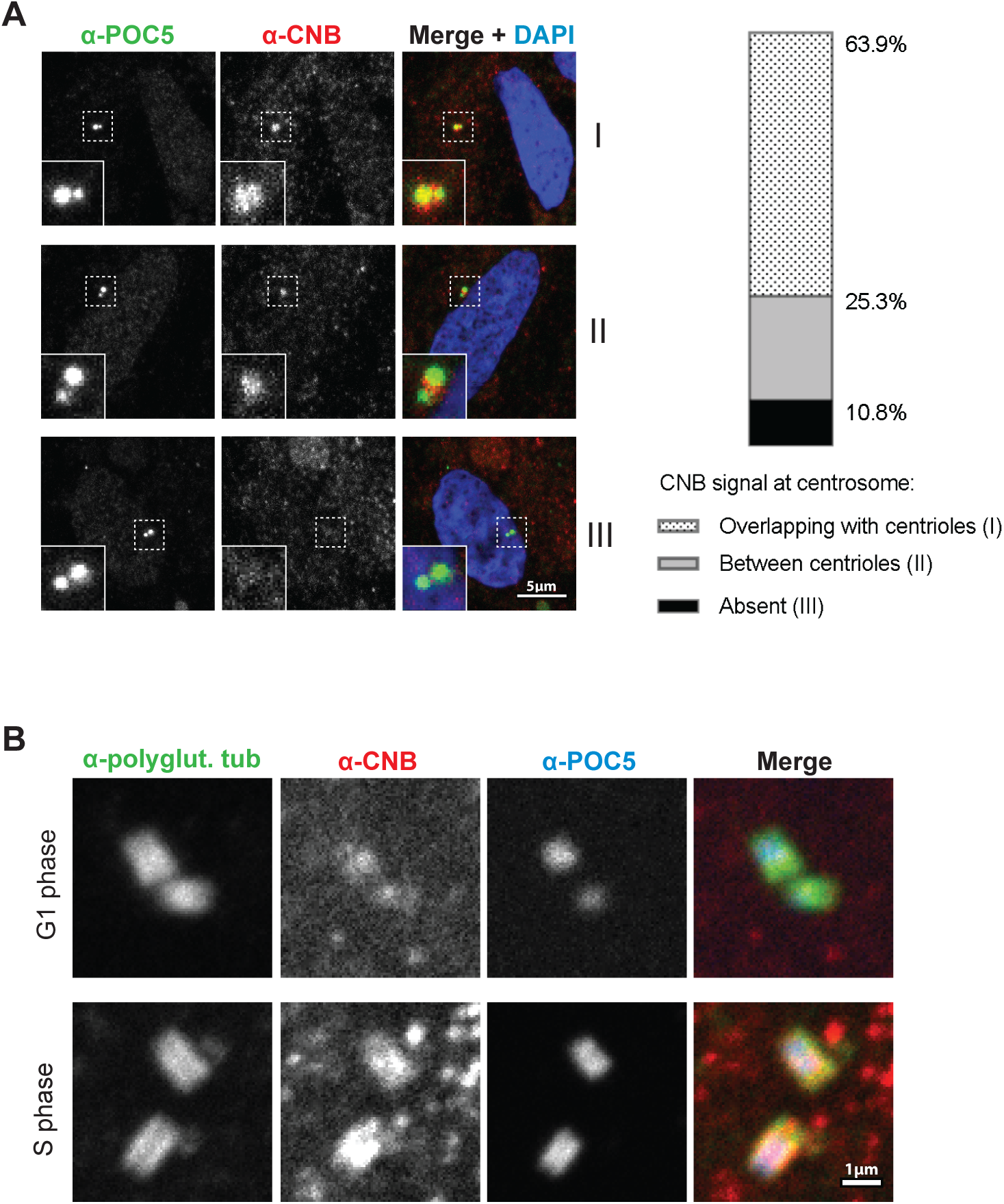
CNB localizes to centrosomes. A. Centrosome associated CNB in cytosol-depleted hTERT-RPE1 cells determined by indirect immunofluorescence. Representative images are maximum projections of confocal z-stacks. Scale bar, 5 μm. Bar chart shows the frequency of different patterns (I, II or III). B. Centrosomal localization of CNB in hTERT-RPE1 cells imaged with expansion microscopy. Top: G1 centrosome. Bottom: S-phase centrosomes. Images are obtained from a single z-plane. Scale bar, 1 μm.

To examine the centrosomal localization of CNB in more detail, we performed ultrastructure expansion microscopy (U-ExM) (Gambarotto et al., 2019), in hTERT-RPE1 cells using the same anti-CNB antiserum used in Fig. 2A. Anti-polyglutamylated tubulin identified the microtubule barrel and anti-POC5 marked the lumen. Despite the presence of cytoplasmic CNB, which increased the background signal, some CNB could be seen to co-localize with tubulin at the centriole barrel (Fig. 2B). No CNB staining was observed at primary cilia in hTERT-RPE1 cells (data not shown).

Our results provide the first evidence that a pool of CN specifically localizes to centrosomes, where it may interact directly with centrosomal proteins.

### CN interacts with centriolar protein POC5 in a PxIxIT-dependent manner

To identify CN binding partners at the centrosome, we focused on POC5, which showed PxIxIT-dependent biotinylation by miniTurbo-CNAα under all conditions (Fig. 1B, Table S1) and contains a predicted PxIxIT motif (Fig. 3A) (Wigington et al., 2020). POC5 is required for centriole maturation and ciliogenesis, and forms a scaffold in the centriole lumen by binding to centrin, POC1B and FAM161A, to stabilize the microtubule barrel (Azimzadeh et al., 2009; Le Guennec et al., 2020). Mutations in POC5 cause adolescent idiopathic scoliosis (Hassan et al., 2019) and retinitis pigmentosa (Weisz Hubshman et al., 2018).

**Fig. 3.**
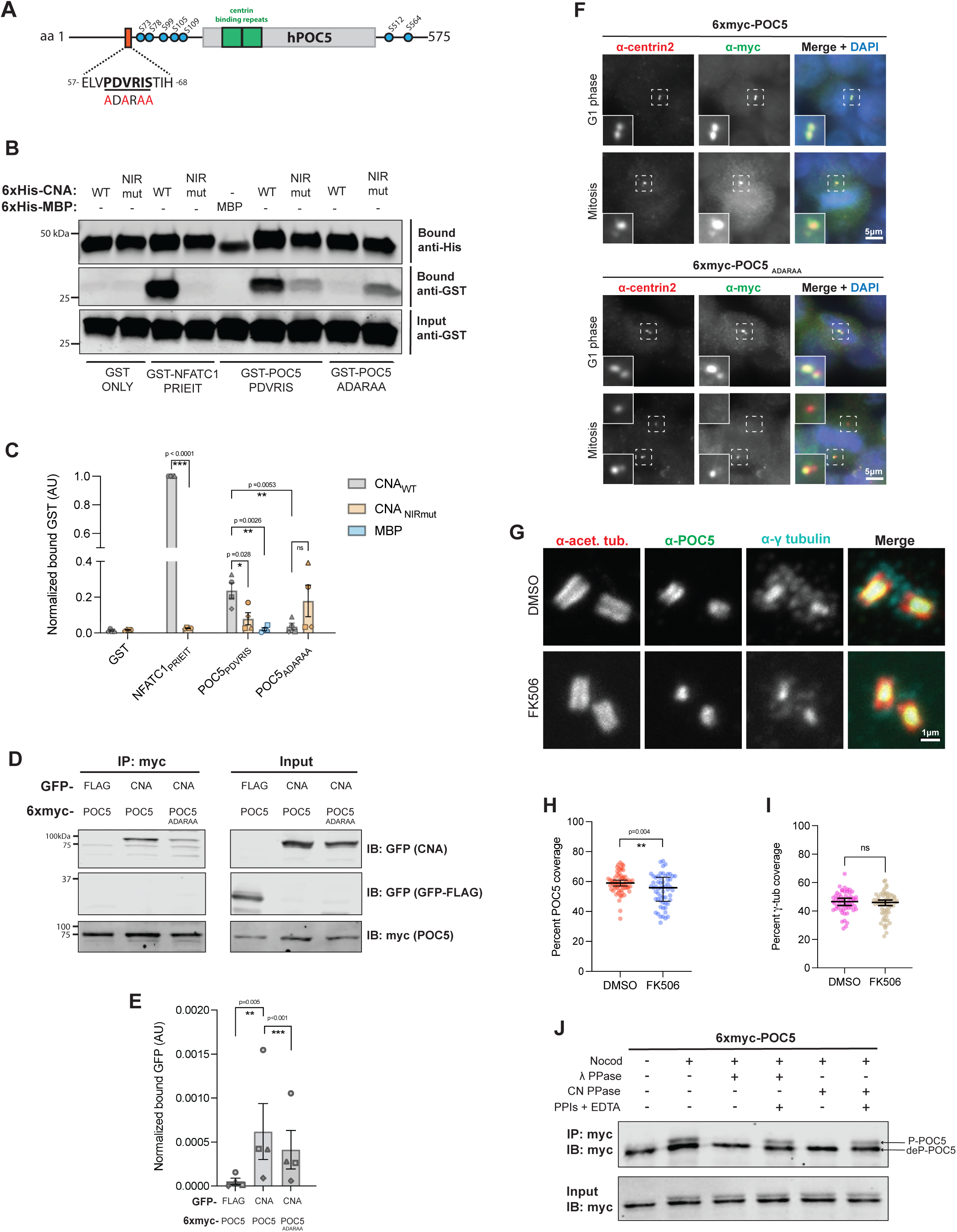
Calcineurin interacts with centriolar protein POC5 in a PxIxIT-dependent manner. A. Schematic of human POC5 showing predicted PxIxIT motif (orange) and mutant ADARAA. Green: centrin binding sequences. Blue circles: phosphorylated residues reported in PhosphoSitePlus (Hornbeck et al., 2015). B. POC5 contains a CN-binding PxIxIT motif. Immunoblot of co-purification of GST-tagged PxIxIT peptides from NFATC1 and POC5 (WT or ADARAAmut) with 6xHis-MBP or CN. C. Quantification of experiment in 3B, i.e., bound GST signal/bound His signal, normalized to input GST. Data are mean ± SEM (n = 4 independent experiments). p-values determined by unpaired, two-tailed t-test. D. CN associates with POC5 in a PxIxIT-dependent manner *in vivo*: Immunoblot of GFP-FLAG or -CNA co-immunoprecipitated with 6xmyc-tagged POC5_WT_ or POC5_ADARAA_ expressed in HeLa cells. E. Quantification of experiment in Fig. 3D shown as bound GFP signal/bound myc signal, normalized to input GFP. Data are mean ± SEM (n = 4 independent experiments). p-values determined by ratio-paired, two-tailed t-test. F. POC5 PxIxIT motif is not required for centriole localization. Immunofluorescence of HeLa cells expressing 6xmyc-POC5, showing POC5 colocalization with centrin. Images obtained from a single z-plane, so only one spindle pole may be in focus in mitosis. The region of interest is magnified in inset. Scale bar, 5 μm. G. Expansion microscopy of hTERT-RPE1 cells treated with DMSO or FK506 for 48 hours, showing POC5 in the centriole lumen and γ-tubulin in the lumen and PCM. Images obtained from a single z-plane. Scale bar, 1 μm. H. CN inhibition decreases luminal distribution of POC5. Median percent of centriole covered by POC5 ± interquartile range (IQR). I. CN inhibition does not alter luminal distribution of γ-tubulin. Median percent of centriole covered by γ-tubulin ± IQR. For H, I: Pooled data from two independent experiments, with 25-30 centrioles analyzed per condition per experiment are shown. p-value > 0.05, determined by unpaired, two-tailed t-test. J. CN dephosphorylates mitotic POC5 *in vitro*. Representative immunoblot of *in vitro* dephosphorylation of 6xmyc-POC5 by 11 phosphatase or CN. Nocod, POC5- expressing cells synchronized with nocodazole. 11 PPase, lambda phosphatase. CN PPase, purified CNA/CNB. PPIs, phosphatase inhibitors. P-POC5, phosphorylated POC5. deP-POC5, dephosphorylated.

To investigate whether CN recognizes POC5’s predicted PxIxIT motif ^57^PDVRIS^68^ (Fig. 3A), we fused WT and mutant POC5 peptides to glutathione S-transferase (GST) and tested their co-purification with 6xHis-CNAα_WT_/CNB, or 6xHis- CNAα_NIRmut_/CNB *in vitro*. CN_WT_ co-purified with the POC5 WT peptide (PDVRIS) and more robustly with a known PxIxIT peptide from NFATC1 (PRIEIT), but not with GST alone. Co-purification of CN_WT_ was significantly reduced with mutant POC5 peptide ADARAA, and CN_NIRmut_ co-purified weakly with all peptides (Figs 3B, C) consistent with PxIxIT-mediated binding.

To examine CN interaction with full-length POC5, we transiently expressed GFP-CNAα_WT_ or GFP-Flag in HEK293T cells together with 6xmyc-POC5_WT_ and -POC5_ADARAA_ and measured the amount of CN that co-immunoprecipitated with POC5. GFP-CNAα_WT_ (but not GFP-FLAG) co-immunoprecipitated with POC5_WT_ and showed reduced interaction with POC5_ADARAA_ (Figs 3D, E), indicating that full-length POC5 binds directly to CN through its identified PxIxIT motif.

Next, we sought to identify a functional role for POC5 binding to CN. To determine whether the POC5 PxIxIT motif is required for its localization to centrioles, we carried out indirect immunofluorescence of HeLa cells transiently expressing 6xmyc-POC5 (POC5_WT_ or mutated PxIxIT POC5_ADARAA_). Both proteins co-localized with centrin at centrioles in G1 and at spindle poles in mitosis (Fig. 3F), and preferentially localized to one centriole, presumably the mother, as described (Azimzadeh et al., 2009). Furthermore, centriole numbers were unaffected in POC5_ADARAA_ and POC5_WT_ -expressing cells (data not shown). Although no CN-dependence was revealed by these analyses, we reasoned that this qualitative determination of POC5 localization under overexpressing conditions might not be sufficiently sensitive to detect CN-dependent regulation of POC5 at centrioles.

Therefore, we used U-ExM to more precisely examine the distribution of endogenously expressed POC5, as well as γ-tubulin, in the centriolar lumen under both control (DMSO) and CN-inhibited (FK506) conditions. POC5 is required for γ-tubulin localization to the centriole lumen, but not to the PCM (Schweizer et al., 2021). Cells were imaged using antisera against POC5, γ-tubulin, and acetylated tubulin, which marks the centriole barrel. We found that POC5 coverage of the centriole lumen was significantly reduced in centrioles from CN-inhibited compared to control cells (Figs 3G-I, S3A). Thus, CN activity may modify POC5 association with or distribution within the centriole lumen, while not disrupting its ability to recruit γ-tubulin.

Together, these findings show that CN interacts directly with POC5 via a PxIxIT motif, and that CN activity affects POC5 distribution at the centriole lumen. However, further analyses are required to determine if CN binding to POC5 is required proper POC5 distribution and/or for CN localization to centrosomes.

### CN dephosphorylates POC5 *in vitro*

We hypothesized that CN activity modifies POC5 distribution by regulating its phosphorylation and set out to test whether POC5 is a CN substrate. POC5 is phosphorylated on several serine residues (Fig. 3A), although the functional significance of these modifications is unknown. POC5 is phosphorylated in mitosis (Azimzadeh et al., 2009); thus, 6xmyc-POC5_WT_ was immunopurified from nocodazole-synchronized, mitotic HeLa cells and incubated *in vitro* with either constitutively active, truncated 6xHis-CNAα_WT_/CNB missing the autoinhibitory tail, or the non-specific 11 phosphatase. POC5 phosphorylation was assessed by visualizing electrophoretic mobility shifts via immunoblotting. POC5 appeared as a doublet with the slower migrating band corresponding to phosphorylated POC5 (p-POC5, Fig. 3J, Input). Treatment with either CN or λ phosphatase eliminated the p-POC5 band, which was preserved by the addition of phosphatase inhibitors (Fig. 3J). Thus, CN dephosphorylates mitotic POC5 *in vitro*.

However, in extracts of mitotic cells expressing 6xmyc-POC5_WT_ or -POC5_ADARAA_, neither CN activation (ionomycin + Ca^2+^), nor inhibition (FK506) altered POC5 electrophoretic migration (Fig. S3B). POC5 is also hyperphosphorylated in nuclear/centrosomal fractions isolated from asynchronous cell cultures (Azimzadeh et al., 2009), but this population of POC5 also failed to show either Ca^2+^ or CN-dependent changes in electrophoretic mobility (Fig. 3SC). Thus, while CN dephosphorylates POC5 *in vitro*, this activity has yet to be demonstrated *in vivo*.

Together, our analyses show that CN interacts directly with POC5 and regulates its distribution within the centriole lumen--possibly by dephosphorylating POC5. While we were unable to demonstrate CN-dependent regulation of POC5 phospho-status i*n vivo*, these analyses await further information about the kinases that target POC5 and the timing and functional significance of these modifications.

### CN inhibition promotes primary cilia elongation

Besides its role in centriole stabilization, POC5 is known to regulate ciliation. Both POC5 and CEP97 (another CN-proximal protein, Fig. 1B) are required for cilia assembly, maintenance, signaling and retraction (Hassan et al., 2019; Schweizer et al., 2021; Spektor et al., 2007) High affinity CN-binding partners CaM, FG-repeat nucleoporins and RCAN2 also localize to basal bodies and regulate ciliation (Kee et al., 2012; Plotnikova et al., 2012; Stevenson et al., 2018), suggesting possible CN-dependent modulation of these processes. To examine whether CN regulates ciliation, IMCD3 cells were grown to near confluency (∼20% ciliated) (Figs 4A, B) and then plated at a high density to induce ciliation via contact inhibition (Joly et al., 2006), while simultaneously treating them with DMSO (control) or the CNI FK506. After 48 hours, ∼50% of either DMSO or FK506-treated cells displayed a cilium (Fig. 4B), indicating that CN activity is not required for ciliogenesis. However, we noticed that the cilia produced were significantly longer in CN-inhibited cells compared to control. Under control conditions, cilia length at 24 hours (median = 6.27 μm) was maintained over the next 24 hours. In contrast, cilia were longer in CN-inhibited cells at 24 hours (median = 7.23 μm) and continued to grow (48 hours, median = 8.59 μm, Fig. 4C).

**Fig. 4.**
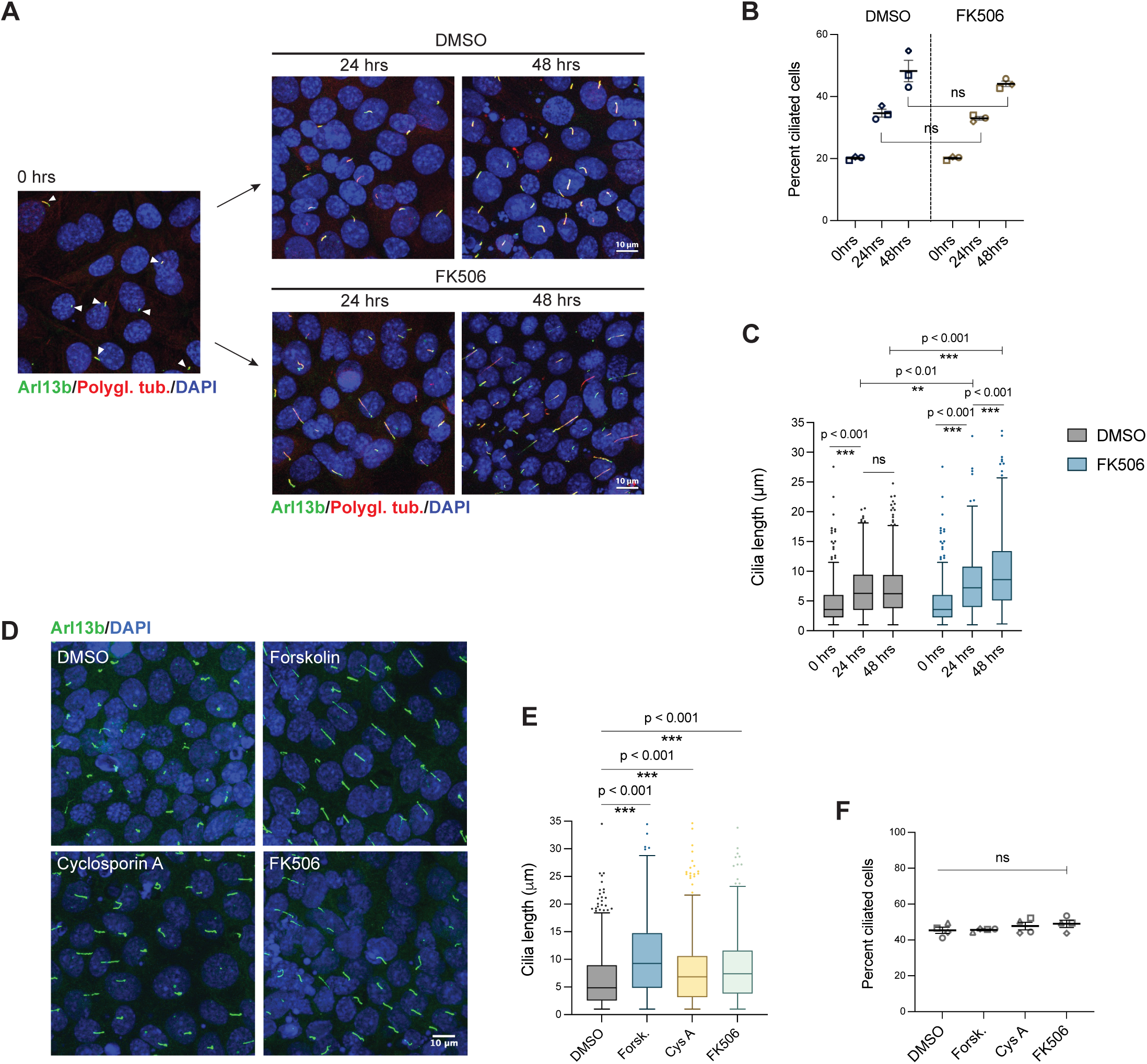
Calcineurin inhibition promotes cilia elongation. A. CN inhibition increases cilia length without disrupting ciliogenesis. Cilia of IMCD3 cells at 0, 24 and 48 hours of treatment with DMSO or FK506. White arrows in the left-most panel point to short cilia. Images are maximum projections of confocal z-stacks. Scale bar, 10 μm. B. Percentage of ciliated IMCD3 cells shown in Fig. 4A. Horizontal lines are mean ± SEM (n = 3 independent experiments). Number of cells analyzed: 0 hrs: n=1319, 24 hrs DMSO: n=2176, 24 hrs FK506: n=2468, 48 hrs DMSO: n=3710, 48 hrs FK506: n=4120. p-values > 0.05, determined by unpaired, two-tailed t-test. C. Length of cilia shown in Fig. 4A. Boxplots: median length ± IQR. Whiskers: median ± 1.5 x IQR. Data pooled from 3 independent experiments. Cilia measured: 0 hrs: n=331, 24 hrs DMSO: n=556, 24 hrs FK506: n=256, 48 hrs DMSO: n=931, 48 hrs FK506: n=429. p-values determined by two-tailed Mann-Whitney test. D. Forskolin and CN inhibitors increase cilia length without altering cilia number. Cilia of IMCD3 cells treated with DMSO, forskolin, or CNIs for 3 hours. Images are maximum projections of confocal z-stacks. Scale bar, 10 μm. E. Length of cilia shown in Fig. 4D. Data shown for one of four replicates; Additional replicates, Fig. 4SA. Cilia measured: DMSO: n=537, forskolin: n=514, cyclosporin A: n=553, FK506: n=638. p-values determined by two-tailed Mann-Whitney test. F. Percentage of ciliated IMCD3 cells shown in Fig 4D. Mean ± SEM shown (n = 4 independent experiments). >1200 cells analyzed per replicate per treatment. p-values > 0.05 determined by unpaired, two-tailed t-test.

Previous studies show that Ca^2+^ and PKA signaling work antagonistically to acutely regulate ciliary length, and that adenylyl cyclase activation by forskolin, or inhibition of Ca^2+^ entry, both lengthen cilia by increasing cAMP levels and PKA activity (Besschetnova et al., 2010). To examine CN’s role in maintaining cilia length, we treated confluent, ciliated IMCD3 cells with CysA or FK506 for 3 hours, which significantly lengthened cilia compared to the DMSO control (Figs 4D-E, 4SA). Elongation was even more dramatic with forskolin (Figs 4D-E, 4SA). However, none of these treatments altered the proportion of ciliated cells (Fig. 4F), suggesting that CN regulates mechanisms maintaining proper cilia length rather than cilium assembly.

In sum, our results reveal that CN localizes to centrosomes, and suggest that CN modifies one or more aspects of centriolar and ciliary homeostasis. Most prior research has focused on centrosome regulation by kinases, but our findings highlight a need to understand phosphatase and Ca^2+^ signaling at these organelles. Ca^2+^ signals have been observed at centrosomes (Helassa et al., 2019), where they activate downstream effectors like CaM (Plotnikova et al., 2012) and centrin. In fact, centrin requires Ca^2+^ to localize to centrosomes and to interact with the structural protein POC5 (Khouj et al., 2019). CN binds directly to POC5 through a PxIxIT motif and alters POC5 distribution within the centriole, suggesting that it may directly regulate this protein despite our inability to detect CN-dependent changes in POC5 phosphorylation *in vivo*. POC5 promotes ciliogenesis by recruiting the augmin complex and the γ-tubulin ring complex (γ-TuRC) to the centriole lumen (Hassan et al., 2019; Schweizer et al., 2021). Of the 8 augmin complex subunits, HAUS5, HAUS6 and HAUS8 have all been identified as CN- proximal with two PxIxIT motifs predicted within HAUS6 (Wigington et al., 2020). Thus, the possibility that CN regulates POC5, augmin or other centriolar components to regulate centriolar stability and/or ciliary function warrants further investigation.

We also discovered that CN regulates cilia length, although the mechanism underlying this effect remains to be determined. Our findings are consistent with previous studies which localized RCAN2, a negative regulator of CN, to centrioles and basal bodies and showed that its depletion resulted in shorter cilia (Stevenson et al., 2018). CN may alter cilia length through effects on POC5 or other structural proteins. Alternatively, CN may mediate documented Ca^2+^-dependent regulation of cAMP/ PKA signaling (Besschetnova et al., 2010), perhaps via binding to AKAP150/AKAP5, a scaffold for CN and PKA that localizes to primary cilia, centrosomes and mitotic spindles. In cilia, AKAP5 associates with adenylyl cyclases AC5/6 and the ciliary Ca^2+^ channel polycystin-2 (PC2) (Choi et al., 2011). Remarkably, the CNA ortholog in *Caenorhabditis elegans,* Tax-6, targets PC2 to cilia (Hu et al., 2006), although this has not been investigated in mammals. Abnormal cilia length is a common phenotype of ciliopathies in highly ciliated organs, such as the kidney and retina (Moreno-Leon et al., 2021; Veland et al., 2009). Thus, elucidating the mechanisms by which CN regulates cilia length and centriolar function promises to improve our current understanding and treatment of ciliary disorders.

## Materials and methods

### Cell lines and culture

Cells were cultured at 37 °C in 5% CO_2_. HEK293T, HeLa and mouse IMCD3 cells were grown in Dulbecco’s Modification of Eagle’s Medium (DMEM) with 4.5g/L glucose, L-glutamine, and sodium pyruvate (Corning, 10-013-CV) supplemented with 10% fetal bovine serum (FBS, Benchmark^TM^, Gemini Bio Products, 100-106). HEK293T and HeLa cells were a gift from Jan Skotheim’s lab at Stanford University. IMCD3 cells were a gift from Peter Jackson’s lab at Stanford University. hTERT-RPE1 cells were cultured in DMEM/F12 media (Gibco^TM^, 11320033) supplemented with 10% FBS. hTERT-RPE1 cells were a gift from Tim Stearns’ lab at Stanford University. Parental HEK293 Flp-In T-REx cells were provided by Anne-Claude Gingras, University of Toronto, and cultured in DMEM supplemented with 10% FBS, 3 μg/mL blasticydin (Research Products International, B12150) and 100 μg/mL zeocin (Gibco™, R25001) prior to stable plasmid integration. Mycoplasma testing was conducted monthly using a mycoplasma PCR detection kit (ABM, G238). Human and mouse cell lines were authenticated by STR profiling (Almeida et al., 2019; Barallon et al., 2010). IMCD3 cells in particular were authenticated using the ATCC Mouse Cell Authentication Service (ATCC, 137-XV).

### MiniTurbo-3xFLAG stable cell lines

MiniTurbo-3xFLAG constructs were generated via Gateway cloning into pDEST 5’ miniTurbo-3xFLAG pcDNA5 FRT TO. Stable cell lines were generated in HEK293 Flp-In T-REx cell pools as described previously for BirA*-FLAG (Hesketh et al., 2017). Doxycycline-inducible, miniTurbo-expressing HEK293 Flp-In T-REx were cultured in antibiotic selection media: DMEM supplemented with 10% FBS, 3 μg/mL blasticydin and 200 μg/mL hygromycin-B (Research Products International, H75000-1). MiniTurbo expression was induced with addition of 1 μg/mL doxycycline (Sigma-Aldrich, D9891) for 48 hours. During BioID assays, stable cell lines were cultured in DMEM supplemented with 10% FBS previously treated to remove residual biotin (method described below), in order to reduce the possibility of non-specific biotinylation.

### Plasmid transfection

For stable cell line generation, HEK293 Flp-In T-Rex cells were co-transfected with pOG44 Flp-Recombinase Expression Vector (Invitrogen™, V600520), and pcDNA5 FRT TO plasmids expressing appropriate miniTurbo gene fusions, using Lipofectamine™ 2000 (Invitrogen™, 11668027), according to the manufacturer’s instructions. All other plasmid transfections were done using jetOPTIMUS® DNA Transfection Reagent (Polyplus), according to the manufacturer’s instructions.

### Biotin depletion of FBS

To remove residual biotin from serum for BioID assays, Streptavidin Sepharose® High Performance beads were used (Cytiva, 17-5113-01). 50 μL of packed bead volume was rinsed three times with 1X phosphate-buffered saline (PBS, pH 7.4, Gibco™ 10010049) in sterile conditions. Beads were spun at 500xg for 1 minute to remove the supernatant, then resuspended in 1X PBS equal to bead volume for 1:1 bead/PBS ratio.

Resuspended beads were added to 50 mL of FBS, and allowed to mix in 4°C for 3 hours. The serum was then spun at 1000xg for 5 minutes to pellet the beads, and the supernatant was filtered through a syringe attached to a 0.45 μm low-bind filter under sterile conditions.

### Cell cycle synchronization coupled to biotinylation

For the asynchronous cell population, miniTurbo-3XFLAG HEK293 Flp-In T-Rex cells were cultured in DMEM containing 10% biotin-depleted FBS, 1 μg/mL doxycycline and DMSO for 48 hours. Immediately prior to cell collection, 50 μM D-biotin (Bio Basic, BB0078) was added to the media for 15 minutes at 37 °C. For G1/S synchronization, cells were cultured in DMEM containing 10% biotin-depleted FBS and 1 μg/mL doxycycline on day 1. On day 2, 2.5 μM thymidine (Millipore-Sigma, T9250) was added to the media for 14 hours at 37 °C. Cultures were then rinsed with 1X PBS and fresh media with 10% biotin-depleted FBS and 1 μg/mL doxycycline were added for 10 hours at 37 °C. 2.5 μM thymidine was then added to the media for 24 hours at 37 °C. Cultures were again rinsed with 1X PBS. Fresh media with 10% biotin-depleted FBS, 1 μg/mL doxycycline and 50 μM D-biotin were added for 15 minutes immediately prior to cell collection. For mitotic synchronization, cells were cultured in DMEM containing 10% biotin-depleted FBS and 1 μg/mL doxycycline on day 1. On day 2, 9 μM RO-3306 (Selleck Chemicals, S7747) was added to the media for 20 hours at 37 °C. Cells were then rinsed with 1X PBS and incubated in DMEM with 10% biotin-depleted FBS and 1 μg/mL doxycycline at 37 °C for 45 minutes. 50 μM D-biotin was then added to the media for 15 minutes, and cells were finally collected one hour post-RO-3306 release. Cells were collected by the addition of warm Trypsin-EDTA Solution, 0.25% (Gibco™, 25200056), and a subset of them were kept resuspended in media for further analysis by flow cytometry and immunoblotting. Collected cells were pelleted by centrifugation at 500xg for 5 minutes. Cell pellets were weighed, frozen in liquid nitrogen and stored at - 80°C until further analysis. Each biotinylation experiment was performed twice, resulting in two biological replicates, or two cell pellets, per condition.

### Validation of BioID samples by flow cytometry and immunoblotting

To prepare for flow cytometry, approximately 1x10^6^ synchronized HEK293 Flp-In T-Rex cells were resuspended in 1 mL 3% paraformaldehyde (PFA) solution and incubated at 37 °C for 10 minutes. Ice-cold methanol (at a volume ratio of 1:9 PFA/methanol) was then added drop-wise to the cell suspension, which was incubated in ice for 30 minutes. Fixed cells were pelleted by centrifugation at 1000xg at 4°C for 10 minutes and the supernatant was removed. The remaining pellet was resuspended in 500 μL bovine serum albumin (BSA, 3% solution in PBS, Sigma-Aldrich, A3294) and centrifuged again at 1000xg at 4°C for 10 minutes. The supernatant was removed, and the pellet was resuspended in 500 µL of DAPI (Cayman Chemical, 14285, 20 μg/mL solution in PBS). The cells were incubated with DAPI at room temperature for 15 minutes, protected from light, and then analyzed for DAPI fluorescence (405 nm laser, VL-1 channel) using the Attune™ NxT Flow Cytometer (Invitrogen), until 50,000 cells had passed through the flow cytometer for each sample. Data was analyzed using the Attune™ NxT Software v3.1.2.

To validate bait expression and successful cell cycle synchronization, synchronized and induced HEK293 Flp-In T-Rex cells were pelleted by centrifugation at 500xg for 5 minutes, frozen in liquid nitrogen, thawed and lysed with RIPA buffer (50 mM Tris-HCl pH 8, 150 mM NaCl, 1% Triton X-100, 0.5% sodium deoxycholate, 0.1% SDS). Samples were resolved by SDS-PAGE and immunoblotted with rabbit anti-β-actin (1:3000, LI-COR Biosciences, 926-42210) as a loading control, mouse anti-FLAG M2 (1:5000, Sigma-Aldrich, F1804) to detect bait expression, rabbit anti-cyclin A2 (1:5000, Abclonal, A2891) as a G1/S marker and rabbit anti-phospho-histone H3 Ser10 (1:500, Millipore, 06-570) as a mitotic marker.

### Immunoblotting

For immunoblotting, samples were denatured with 2X or 6X sodium dodecyl sulfate (SDS) Laemmli buffer and boiled at 95°C for 5 minutes. Protein concentrations were determined using the Pierce™ BCA Protein Assay Kit (Thermo Fisher Scientific, 23225), according to the manufacturer’s instructions. Equal amounts of protein (20-40 μg) were separated by SDS-PAGE. Proteins were transferred to a nitrocellulose membrane (Bio-Rad, 162-0112). The membrane was blocked with SuperBlock blocking buffer (Fisher, PI37535) at room temperature for 30 minutes and then incubated with primary antibodies at 4°C overnight, or at room temperature for 1 hour. Next, the membrane was incubated with one or both of the following secondary antibodies: IRDye 680RD Goat anti-mouse IgG (H + L) (1:15,000, Li-COR Biosciences 926-68071) and IRDye 800CW Goat anti-rabbit IgG (H + L) (1:15,000, Li-COR Biosciences 926-32211) at room temperature for 1 hour. All blots were imaged with the Li-COR Odyssey imaging system and analyzed using Image Studio (Li-COR Biosciences).

### Biotin-streptavidin affinity purification and on-bead trypsin digest

Frozen cell pellets were first thawed and then lysed, bound to streptavidin-sepharose beads, trypsinized, dried, and prepared for analysis by mass spectrometry exactly as described in the protocol detailed in section 3.4.1 in Hesketh et al., 2017.

### Mass spectrometry data acquisition

Both data-dependent acquisition (DDA) and data-independent acquisition (DIA) were performed. For DDA, LC-MS/MS, affinity purified and digested peptides were analyzed using a nano-HPLC (High-performance liquid chromatography) coupled to MS. One-quarter of the sample was used. Nano-spray emitters were generated from fused silica capillary tubing, with 100 µm internal diameter, 365 µm outer diameter and 5-8 µm tip opening, using a laser puller (Sutter Instrument Co., model P-2000, with parameters set as heat: 280, FIL = 0, VEL = 18, DEL = 2000). Nano-spray emitters were packed with C18 reversed-phase material (Reprosil-Pur 120 C18-AQ, 3 µm) resuspended in methanol using a pressure injection cell. Sample in 5% formic acid was directly loaded at 800 nl/min for 20 minutes onto a 100 µm x 15 cm nano-spray emitter. Peptides were eluted from the column with an acetonitrile gradient generated by an Eksigent ekspert™ nanoLC 425, and analyzed on a TripleTOF™ 6600 instrument (AB SCIEX, Concord, Ontario, Canada). The gradient was delivered at 400 nl/min from 2% acetonitrile with 0.1% formic acid to 35% acetonitrile with 0.1% formic acid using a linear gradient of 90 min. This was followed by a 15 minute wash with 80% acetonitrile with 0.1% formic acid, and equilibration for another 15 minutes to 2% acetonitrile with 0.1% formic acid. The total DDA protocol is 135 minutes. The first DDA scan had an accumulation time of 250 ms within a mass range of 400-1800 Da. This was followed by 10 MS/MS scans of the top 10 peptides identified in the first DDA scan, with accumulation time of 100 ms for each MS/MS scan. Each candidate ion was required to have a charge state from 2-5 and a minimum threshold of 300 counts per second, isolated using a window of 50 mDa. Previously analyzed candidate ions were dynamically excluded for 7 seconds. For DIA, LC-MS/MS, affinity purified and digested peptides were analyzed using a nano-HPLC (High-performance liquid chromatography) coupled to MS. One-quarter of the sample was used. Nano-spray emitters were generated from fused silica capillary tubing, with 100 µm internal diameter, 365 µm outer diameter and 5-8 µm tip opening, using a laser puller (Sutter Instrument Co., model P-2000, with parameters set as heat: 280, FIL = 0, VEL = 18, DEL = 2000). Nano-spray emitters were packed with C18 reversed-phase material (Reprosil-Pur 120 C18-AQ, 3 µm) resuspended in methanol using a pressure injection cell. Sample in 5% formic acid was directly loaded at 800 nl/min for 20 minutes onto a 100 µm x 15 cm nano-spray emitter. Peptides were eluted from the column with an acetonitrile gradient generated by an Eksigent ekspert™ nanoLC 425, and analyzed on a TripleTOF™ 6600 instrument (AB SCIEX, Concord, Ontario, Canada). The gradient was delivered at 400 nl/min from 2% acetonitrile with 0.1% formic acid to 35% acetonitrile with 0.1% formic acid using a linear gradient of 90 minutes. This was followed by a 15 minute wash with 80% acetonitrile with 0.1% formic acid, and equilibration for another 15 minutes to 2% acetonitrile with 0.1% formic acid. The total DIA protocol is 135 minutes. The first DIA scan had an accumulation time of 250 ms within a mass range of 400-1800 Da. This was followed by 54 MS/MS scans with differing mass windows, with an accumulation time of 65 ms per scan.

### Mass spectrometry data analysis

Mass spectrometry data generated were stored, searched and analyzed using ProHits laboratory information management system (LIMS) platform (Liu et al., 2016). Within ProHits, WIFF files were converted to an MGF format using the WIFF2MGF converter and to an mzML format using ProteoWizard (V3.0.10702) and the AB SCIEX MS Data Converter (V1.3 beta).

DDA acquisition data was searched using Mascot (V2.3.02) (Perkins et al., 1999) and Comet (V2016.01 rev.2, (Eng et al., 2013).The spectra were searched with the human and adenovirus sequences in the RefSeq database (version 57, January 30th, 2013) acquired from NCBI, supplemented with “common contaminants” from the Max Planck Institute (http://www.coxdocs.org/doku.php?id=maxquant:start_downloads.htm) and the Global Proteome Machine (GPM; ftp://ftp.thegpm.org/fasta/cRAP), forward and reverse sequences (labeled “gi|9999” or “DECOY”), sequence tags (BirA, GST26, mCherry and GFP) and streptavidin, for a total of 72,481 entries. Database parameters were set to search for tryptic cleavages, allowing up to 2 missed cleavages sites per peptide with a mass tolerance of 35 ppm for precursors with charges of 2+ to 4+ and a tolerance of 0.15 amu for fragment ions. Variable modifications were selected for deamidated asparagine and glutamine and oxidized methionine. Results from each search engine were analyzed through TPP (the Trans-Proteomic Pipeline, v.4.7 POLAR VORTEX rev 1) via the iProphet pipeline (Shteynberg et al., 2011).

DIA acquisition data was searched using MS-GFDB (Wang et al., 2015). The spectra were searched with the human and adenovirus sequences in the RefSeq database (version 57, January 30th, 2013) acquired from NCBI, supplemented with “common contaminants” from the Max Planck Institute (http://www.coxdocs.org/doku.php?id=maxquant:start_downloads.htm) and the Global Proteome Machine (GPM; ftp://ftp.thegpm.org/fasta/cRAP), for a total of 36,361 entries. Database parameters were set to search for tryptic cleavages, with a mass tolerance of 50 ppm for precursors with charges of 2+ to 4+ and a peptide length of 8 to 30 amino acids. Oxidized methionine was set as the variable modification. DDA files were used to generate a spectral library for the SWATH files.

### *SAINT* analysis

The SAINT analysis tool is used to identify high-confidence protein interactors versus control samples (Teo et al., 2014). SAINTexpress version 3.6.1 was used for DDA and version 3.6.3 for MSPLIT. SAINT analysis was performed using two biological replicates per bait for both DDA and MSPLIT. Six negative control experiments with miniTurbo-3xFLAG-alone samples were conducted for BioID; two asynchronous replicates, two in G1/S and two in mitosis. SAINT probabilities were calculated independently for each sample, averaged (AvgP) across biological replicates and reported as the final SAINT score. Prior to applying SAINT, proteins were filtered with iProphet ≥ 0.95 and unique peptides ≥ 2 for DDA and unique peptides ≥ 2 for MSPLIT. Proteins with a BFDR (Bayesian False Discovery Rate) ≤ 0.01 are considered high-confidence protein interactors. Heat maps were generated from SAINT output via ProHits-viz (Knight et al., 2015).

### Filtered prey dataset

For each prey, the averaged spectral counts obtained with miniTurbo alone (2 replicates x 3 conditions) were subtracted from averaged spectral counts with miniTurbo-CNA (wild type or mutant) for each condition (background subtracted spectral counts, Table S1). Final dataset of 41 proteins resulted from 38 preys from DDA analysis and 3 additional preys from DIA/mSPLIT analysis, that were significantly biotinylated by miniTurbo-CNA_WT_ (unique peptides ≥ 2, BFDR ≤ 0.01) in at least one condition. PxIxIT-dependent proteins displayed Log_2_ (spectral counts with miniTurbo-CNA_WT_/miniTurbo- CNA_NIRmut_) ≥ 0.5 for at least one condition. For preys with spectral counts with miniTurbo-CNA_NIRmut_ = 0, values were converted to 0.5 to calculate CNA_WT_/CNA_NIRmut_ ratios.

### GST-PxIxIT peptide purification

Oligos coding for 16-mer peptides with PxIxIT motifs in the center were fused to GST in a pGEX-4T-3 vector and expressed in BL21 (DE3) chemically competent E. coli (Sigma-Aldrich, CMC0014). Bacteria were grown in 37°C until mid-log phase and expression was induced with 1 mM isopropyl-β-D-thiogalactopyranoside (IPTG, P212121, GB-I0920) addition for 2 hours. Bacteria were lysed with CelLytic™ B Cell Lysis Reagent (Millipore-Sigma, B7435) according to the manufacturer’s instructions. Cell lysates expressing GST-peptides were incubated with Glutathione Sepharose™ 4B beads (Cytiva, 17-0756-01) in 4°C for 2-4 hours and the beads were then isolated and eluted through a Bio-Spin® Chromatography Column (Bio-Rad, #7326008) with elution buffer (5 0mM Tris-HCl pH 8, 300 mM NaCl, 0.1% NP-40, 5 mM Dithiothreitol-DTT, 40 mM glutathione, NaOH added to adjust buffer pH to 8). The eluates were allowed to dialyze in 4°C overnight in dialysis buffer (100 mM Tris-HCl pH 8, 150 mM NaCl, 1 mM β-mercaptoethanol) to remove residual glutathione. Purified peptides were stored in 10% glycerol at -80°C.

### 6xHis-tagged calcineurin and maltose binding protein purification

6xHis-tagged human calcineurin A (α isoform, truncated at residue 392), WT or ^330^NIR^332^-^330^AAA^332^ mutant were expressed in tandem with the calcineurin B subunit in a p11 vector in BL21 (DE3) chemically competent *E. coli*. Similarly, 6xHis-tagged, WT maltose binding protein (MBP) was expressed in a p11 vector in BL21 (DE3) chemically competent *E. coli*. Bacteria were grown in 37°C until mid-log phase and expression was induced with 1 mM IPTG at 16°C for 18 hours. Cells were pelleted, washed, and frozen at −80°C for at least 12 hours. Thawed cell pellets were resuspended in lysis buffer (50 mM Tris-HCl pH 7.5, 150 mM NaCl, 0.1% Tween 20, 1 mM β-mercaptoethanol, protease inhibitors) and lysed by sonication using four 1- minute pulses at 40% output. Extracts were clarified using two rounds of centrifugation (20,000xg, 20 minutes, 4°C) and then bound to Ni-NTA agarose beads (Invitrogen, R901-15) in lysis buffer containing 5mM imidazole for 2–4 hours at 4°C. Bound beads were loaded onto a Bio Spin® Chromatography Column and washed with lysis buffer containing 20mM imidazole (Sigma-Aldrich, I0250) and eluted with lysis buffer containing 300mM imidazole, pH 7.5. Purified proteins were dialyzed in buffer (50 mM Tris-HCl pH 7.5, 150mM NaCl, 1 mM β-mercaptoethanol) and stored in 10% glycerol at −80°C.

### *In vitro* GST peptide binding

1-2 μg of purified 6xHis-tagged calcineurin or 6xHis-MBP as a negative control was first bound to 10 μL magnetic Dynabeads™ His-Tag Isolation and Pulldown Beads (Thermo Fisher Sci., 10104D) in 490 μL base buffer (50 mM Tris-HCl pH 7.5,150 mM NaCl, 0.1% Tween 20, 1 mM β-mercaptoethanol, protease inhibitors), supplemented with 15 mM imidazole and 0.5 mg/ml BSA for 1.5 hours at 4°C. 7-10 μg of appropriate purified GST-peptides were then added to the binding reaction and incubated further for 2 hours at 4°C. 3% of the total reaction mix was removed as ‘input’ prior to the incubation, boiled in Laemmli sample buffer, and stored at −20°C. The beads were washed in base buffer containing 20mM imidazole and bound proteins were eluted by boiling in Laemmli sample buffer for 5 minutes, followed by SDS–PAGE and immunoblotting with goat anti-GST (1:3000, Cytiva, 27-4577-01) and mouse anti-6xHis (1:3000, Takara Bio, 631212) antibodies. Secondary antibodies used were IRDye 680RD Goat anti-mouse IgG (H + L) (1:15,000, Li-COR Biosciences 926-68071) and IRDye 800CW Donkey anti-goat IgG (H + L) (1:15,000, Li-COR Biosciences 926-32214). GST peptides co-purifying with 6xHis-tagged proteins were normalized to their respective input and amount of His-protein pulled down. Co-purification with CN was reported relative to that of the peptide with the known PxIxIT motif from NFATC1: PALESPRIEITSCLGL. POC5 peptides used were POC5 PDVRIS: KGELVPDVRISTIHDI and POC5 ADARAA Mut: KGELVADARAATIHDI. Statistical significance was determined with unpaired, two-tailed Student’s t test, using GraphPad Prism 9. *In vitro* GST pulldown experiments were performed in three biological replicates.

### Calcineurin and POC5 co-immunoprecipitation assays

HEK293T cells transfected with plasmids expressing 6xmyc-tagged POC5 (WT or ^57^ADARAA^68^ mutant) and GFP-tagged FLAG or GFP-CNA subunit α isoform were washed with 1X PBS and harvested. Cell pellets were snap-frozen in liquid nitrogen and stored at −80°C until use. Thawed cell pellets were lysed with lysis buffer (50 mM Tris-HCl pH 7.5, 150 mM NaCl, 1% NP-40), supplemented with Halt™ Protease and Phosphatase Inhibitor Cocktail (Thermo Scientific, 78440) and subjected to fine needle aspiration through a sterile 27.5-gauge needle. Cell lysates were clarified by centrifugation at 16,000xg for 20 minutes in 4°C and protein concentrations were determined using the Pierce™ BCA Protein Assay Kit, according to the manufacturer’s instructions. 1 mg of protein from each lysate was added to 20 μL of pre-washed anti-c-myc magnetic beads (Med Chem Express, HY-K0206) and the volume of each reaction was equalized to 500 μL with binding buffer (50 mM Tris-HCl pH 7.5, 150 mM NaCl, 0.5% NP-40, Halt™ Protease and Phosphatase Inhibitor Cocktail). 2.5% of the total reaction mix was removed as ‘input’ prior to the incubation, boiled in Laemmli sample buffer, and stored at −20°C. The reactions were then rotated gently at 4°C overnight. Beads were washed four times with wash buffer (0.5% Triton X-100, Halt™ Protease and Phosphatase Inhibitor Cocktail in 1X PBS) and boiled in 2X Laemmli sample buffer for 5 minutes. 50% of ‘input’ and ‘immunoprecipitated’ fractions were resolved by SDS-PAGE and immunoblotted with rabbit anti-myc (1:3000, 71D10, Cell Signaling Technology, 2278S) and mouse anti-GFP (1:3000, Santa Cruz Biotechnology, sc-9996) antibodies. Four biological replicates of this experiment were performed. GFP-tagged proteins co-immunoprecipitating with 6xmyc-tagged POC5 were normalized to their respective input and then over the amount of 6xmyc-POC5 bound to the beads. Statistical significance was determined with ratio-paired, two-tailed Student’s t test, using GraphPad Prism 9.

### Immunoprecipitation and *in vitro* dephosphorylation of POC5

HeLa cells were transfected with plasmid expressing wild-type 6xmyc-tagged POC5 and divided between two plates: one where cells were treated with DMSO, and another where cells were treated with 100 ng/mL nocodazole (Cell Signaling Technology, 2190S) for 18 hours at 37°C. DMSO or nocodazole were washed out with 1X PBS and cells were incubated with fresh, drug-free media for an hour at 37°C. Cells were harvested, pelleted, frozen in liquid nitrogen and stored at −80°C until use. Thawed cell pellets were lysed with lysis buffer (50mM Tris-HCl pH 7.5, 150mM NaCl, 1% NP-40), supplemented with Halt™ Protease and Phosphatase Inhibitor Cocktail and subjected to fine needle aspiration through a sterile 27.5-gauge needle. Cell lysates were clarified by centrifugation at 16,000xg for 20 minutes in 4°C and protein concentrations were determined using the Pierce™ BCA Protein Assay Kit, according to the manufacturer’s instructions. 500 μg of protein from each lysate was added to 20 μL of pre-washed anti-c-myc magnetic beads (Med Chem Express, HY-K0206) and the volume of each reaction was equalized to 500 μL with binding buffer (50 mM Tris-HCl pH 7.5, 150 mM NaCl, 0.5% NP-40, Halt™ Protease and Phosphatase Inhibitor Cocktail). 2.5% of the total reaction mix was removed as ‘input’ prior to the 2 hour incubation, boiled in 2X Laemmli sample buffer, and stored at 4°C. The bead binding mixtures were then rotated gently at 4°C for 2 hours. Beads were washed twice with wash buffer (0.5% Triton X-100, Halt™ Protease and Phosphatase Inhibitor Cocktail in 1X PBS) and then washed twice with either λ dephosphorylation buffer (1 mM MnCl_2,_ 1X PMP buffer, New England Biolabs, P0753, protease inhibitors) or CN dephosphorylation buffer (50 mM Tris-HCl pH 8, 100 mM NaCl, 6 mM MgCl_2,_ 1 mM CaCl_2,_ 1mM DTT, protease inhibitors). Finally, the beads were incubated in λ dephosphorylation buffer or CN dephosphorylation buffer, with or without phosphatase addition (0.25 μL of λ phosphatase in 50 μL reaction volume, New England Biolabs, P0753 or 200 nM purified 6xHis-Calcineurin) and with or without phosphatase inhibitor addition (Halt™ Protease and Phosphatase Inhibitor Cocktail and 5 mM EDTA), as required. Dephosphorylation was allowed to occur for 45 minutes at 30°C under constant shaking. Reactions were stopped with 2X Laemmli buffer and boiled for 5 minutes. Proteins were analyzed on 6% acrylamide SDS-PAGE gels followed by immunoblotting with rabbit anti-myc (1:3000, 71D10, Cell Signaling Technology, 2278S). *In vitro* dephosphorylation experiments were performed in three biological replicates.

### Cytosol extraction and immunofluorescence microscopy

HeLa, IMCD3 or hTERT-RPE1 cells were grown on poly-L-lysine (Sigma-Aldrich, P4707) pre-treated 12 mm, #1.5H glass coverslips (ThorLabs). If cytosol extraction was not required, cells were directly fixed in ice-cold methanol at -20°C for 15 minutes. For cytosol-extracted RPE1 cells, the coverslips were first washed with warm 0.02% digitonin in PBS solution and rocked gently for 5 minutes at room temperature, followed by one wash with 1X PBS and methanol fixation at -20°C for 15 minutes. Following methanol fixation, coverslips were washed thrice with 1X PBS and then placed in a dark humid chamber and treated with blocking buffer (0.2 M glycine, 0.1% Triton X-100, 2.5% FBS in PBS) for 30 minutes. Coverslips were incubated with primary antibodies diluted in block buffer for 1 hour, washed multiple times with 1X PBS, followed by incubation with secondary antibodies for 45 minutes, multiple washes with 1X PBS and a 15-minute incubation with 5 μg/mL DAPI at room temperature. Coverslips were washed again and mounted on glass slides using ProLong® Diamond Antifade Mountant (Thermo Fisher, P36965).

Primary antibodies used in immunofluorescence: mouse anti-centrin2, clone 20H5 (1:500, EMD Millipore, 04-1624), mouse anti-centrin3 3E6 (1:500, Abnova, H00001070-M01), rabbit anti-myc 71D10 (1:300, Cell Signaling Technology, 2278S), rabbit anti-POC5 CE037 (1:100, Invitrogen, PA524308), mouse anti-CNB (1:100, Sigma-Aldrich, C0581), rabbit anti-calcineurin pan A (1:100, EMD Millipore, 07-1491). Secondary antibodies used: goat anti-mouse Alexa Fluor 594 (1:1000, Invitrogen A11032), goat anti-rabbit Alexa Fluor 488 (1:1000, Invitrogen A11008). Imaging was performed at the Stanford University Cell Sciences Imaging Facility (CCIF RRID:SCR_017787), using an Inverted Zeiss LSM 880 confocal laser scanning microscope with either a 1.3 NA 40x EC Plan Neo oil immersion objective or a 1.4 NA 63x Plan Apo oil immersion objective. Lasers used were Diode 405nm 0.2-0.8mW, HeNe 594nm 2mW and Ar 488nm 25mW. Images were acquired at constant exposure settings within experiments using the Zen Black software (Carl Zeiss). Image J was used for image analysis, line intensity plots and quantification of fluorescence intensities.

### Ultrastructure expansion microscopy (U-ExM)

hTERT-RPE1 cells were grown on 12 mm, #1.5H glass coverslips and treated either with DMSO (control) or 2.5 μM FK506 (LC laboratories, F4900) for 48 hours. Coverslips were fixed in ice-cold methanol at -20°C for 10 minutes and washed with 1X PBS. Following fixation, U-ExM was performed as previously described in (Gambarotto et al., 2019). Briefly, coverslips were incubated overnight in an acrylamide/formaldehyde solution (AA/FA, 0.7% formaldehyde, 1% acrylamide in PBS) at 37°C. Gelation was allowed to proceed in monomer solution (19% sodium acrylate, 10% acrylamide, 0.1% bis-acrylamide, 0.5% ammonium persulfate-APS, 0.5% TEMED) and the coverslips were discarded. Gels were boiled in denaturation buffer (200 mM SDS, 200 mM NaCl, 50 mM Tris pH 9) at 95°C for 1 hour. After denaturation buffer was removed, gels were washed with multiple water rinses and allowed to expand in water at room temperature overnight. Small circles of each expanded gel (approximately 5 mm in diameter) were excised and incubated with primary antibodies diluted in PBSBT buffer (3% BSA, 0.1% Triton X-100 in PBS) on a nutator at 4°C overnight. The next day, gels were washed thrice with PBSBT buffer and incubated with secondary antibodies and 5 μg/mL DAPI diluted in PBSBT, protected from light, on a nutator at 4°C overnight. Immunostained gels were washed once with 1X PBS and thrice with water, and placed in a glass-bottom, poly-L-lysine treated 35mm plate for imaging. Primary antibodies used in expansion microscopy were: mouse anti-polyglutamylated tubulin GT335 (1:500, AdipoGen, AG20B0020C100), rabbit anti-calcineurin pan A (1:100, EMD Millipore, 07-1491), mouse anti-CNB (1:100, Sigma-Aldrich, C0581), rabbit anti-POC5 CE037 (1:500, Invitrogen, PA524308), mouse anti-acetylated tubulin clone 6-11B-1 (1:500, Sigma-Aldrich, T6793), mouse anti-gamma tubulin GTU-88 (1:500, Abcam, ab11316). Secondary antibodies used were: goat anti-mouse IgG1 Alexa Fluor 488 (1:1000, Invitrogen, A21121), goat anti-mouse IgG2b Alexa Fluor 488 (1:1000, Invitrogen, A-21141), goat anti-rabbit Alexa Fluor 647 (1:500, Invitrogen, A21245), goat anti-mouse IgG2b Alexa Fluor 647 (1:500, Invitrogen, A21242), goat anti-mouse IgG2b Alexa Fluor 568 (1:500, Invitrogen, A21144), goat anti-mouse IgG2a Alexa Fluor 568 (1:500, Invitrogen, A21134), goat anti-rabbit Alexa Fluor 568 (1:500, Invitrogen, A11011). All expansion microscopy images were acquired as single planes or Z-stacks collected at 0.27-μm intervals using a confocal Zeiss Axio Observer microscope (Carl Zeiss) with a PlanApoChromat 1.4 NA 63x oil immersion objective, a Yokogawa CSU-W1 head, and a Photometrics Prime BSI express CMOS camera. Slidebook software (Intelligent Imaging Innovations, 3i) was used to control the microscope system. Image J was used for image analysis and quantification of fluorescence intensities.

### Sucrose fractionation for nuclear/centrosomal fractions

HeLa cells were resuspended in 1X PBS buffer, followed by addition of lysis buffer (10 mM Tris-HCl, pH 7.5, 10 mM NaCl, 3 mM MgCl_2_, 1% NP-40, 10% sucrose and protease inhibitors) at 1 mL per 1x10^7^ cells. Lysed cells were centrifuged at 100xg at 4°C for 5 minutes. A small fraction of the supernatant was flash-frozen and stored in -80°C as the cytosolic fraction and the rest was discarded. The pellet containing the nuclei and centrosomes was incubated at room temperature for 30 minutes in the same volume of digestion buffer (10 mM K-PIPES pH 6.8, 50 mM NaCl, 3 mM MgCl_2_, 1 mM EGTA, 10% sucrose, 0.1 mg/mL DNAse I, 0.1 mg/mL RNAse A) as the volume used for lysis buffer. (NH_4_)_2_SO_4_ and NaCl solutions were then added at final concentrations of 0.25 M and 1 M respectively. The mixture was centrifuged at 15,000 xg for 15 minutes over a 1 mL cushion made of 60% sucrose in digestion buffer. After centrifugation, 2 mL of material (1 mL of supernatant and 1 mL of cushion) was collected at the 10-60% sucrose interphase and contents were vortexed, flash-frozen and stored at -80°C.

### Anti-CNB antibody blocking

60 μg of BSA protein (Sigma-Aldrich, A3294) and 60 μg purified 6xHis-CNAα copurified with regulatory subunit CNB were resolved by SDS-PAGE and transferred to a nitrocellulose membrane (Bio-Rad, 162-0112). The membrane was stained with Ponceau S solution (Sigma-Aldrich, P7170) for 15 minutes and rinsed twice with distilled water to visualize the bands of interest. The membrane was excised around the bands corresponding to BSA and CNB and the membrane strips were first washed with water to remove excess Ponceau, and then blocked with 5% milk in TBST buffer (Tris-buffered saline with 0.1% Tween-20) for 1 hour at room temperature. The strips were then rinsed twice in TBST buffer, and twice in PBSBT buffer. Finally, each strip was incubated with 99 μL PBSBT and 1 μL mouse anti-CNB antibody (Sigma-Aldrich, C0581, 1:100), on the nutator for five hours at room temperature. The membrane strips were then discarded and the antibody mixture was added onto coverslips to stain them for immunofluorescence as described.

### Centriole length and percent POC5 / y-tubulin coverage measurements

All measurements were performed on Image J using expansion microscopy, single z-plane images of hTERT-RPE1 centrioles after 48-hour treatment with DMSO or 2.5 μM FK506. Overall centriole length was quantified using acetylated tubulin staining and coverage length was determined by anti-POC5 or anti-γ-tubulin staining, exactly as previously described for GCP4 coverage (Schweizer et al., 2021).

### Percent ciliation and cilia length measurements

To measure changes in the maintenance of existing cilia, IMCD3 cells were grown to confluency on 12 mm, #1.5H glass coverslips. Two days after reaching confluency, cells were treated with DMSO, 10 μM forskolin (Sigma-Aldrich, F3917), 2.5 μM FK506 or 2.5 μM cyclosporin A (Sigma-Aldrich, 30024) for 3 hours at 37°C. Coverslips were fixed in 4% PFA for 10 minutes in room temperature and then prepared for immunofluorescence. Four biological replicates of this experiment were performed. To measure changes during active ciliogenesis, IMCD3 cells were grown to 90% confluency on a 10 cm tissue culture plate containing two 12 mm, #1.5H glass coverslips. The coverslips were removed and fixed in 4% PFA for 10 minutes in room temperature and then prepared for immunofluorescence. The remaining cells on the plate were trypsinized and split into two fully confluent 3.5 cm tissue culture plates with glass coverslips. One plate contained media supplemented with DMSO and the other with 2.5 μM FK506. 24 hours after drug treatment, two coverslips from the DMSO-treated plate and two from the FK506-treated plate were removed, fixed and stained for immunofluorescence. The same procedure was performed again 48 hours after drug treatment began. This experiment was performed in triplicate. Primary antibodies used to stain cilia were rabbit anti-Arl13b (1:250, Proteintech, 17711-1-AP) and mouse anti-polyglutamylated tubulin GT335 (1:500, AdipoGen, AG20B0020C100).

Immunofluorescence and image acquisition was performed as detailed in the ‘cytosol extraction and immunofluorescence’ section of the Methods. Image J was used to count the number of cells (determined from DAPI-stained nuclei in maximum projection of all z-planes) in each image. The CiliaQ plugin for Image J (https://github.com/hansenjn/CiliaQ) (Hansen et al., 2021) with the CANNY 3D segmentation method was used to determine the number and lengths of Arl13b-stained cilia from z-stacks collected at 0.5 μm intervals in the 488 nm channel. Using R Studio, cilia length data were accumulated and filtered so that only single-branched cilia with length over 1 μm were selected for further analysis.

Percent ciliation was determined by #cilia with length > 1 μm (Arl13b as a 3D vector in a z-stack) / #nuclei (DAPI in maximum projection images of a confocal z-stack).

### Statistics

All significance testing was conducted using GraphPad Prism 9. n.s., not significant, *p<0.05, **p<0.01, ***p<0.001.

## Acknowledgements

We thank all members of the Cyert Lab for providing support and editorial assistance; members of the Tim Stearns and Jan Skotheim Labs (Stanford University, CA, USA) for feedback, support, reagents and cell lines, and in particular Mardo Kõivomägi for reagents and expertise regarding phosphorylation assays; Richard Lewis and Kang Shen (Stanford University, CA, USA) for critical feedback; Jan Niklaus Hansen and Dagmar Wachten (University of Bonn, Bonn, Germany) for sharing and providing assistance with the CiliaQ software; Peter Jackson (Stanford University, CA, USA) for IMCD3 cell lines; Anne-Marie Tassin (Institute for Integrative Biology of the Cell, Gif-sur-Yvette, France) and Juliette Azimzadeh (Institut Jacques Monod, Paris, France) for sharing centrosomal fractionation protocols; Kitty Lee and the Stanford University Cell Sciences Imaging Facility (CCIF RRID:SCR_017787) for providing help, training and equipment for confocal imaging.

## Author contributions

Conceptualization: M.S.C., E.T., J.T.W.; Investigation: E.T., J.T.W., C.J.W., I.U.-T.,T.S., A.-C.G.; Data curation: E.T., C.J.W., A.-C.G., I.U.-T.; Writing - original draft: E.T.; Writing - review & editing: M.S.C., I.U.-T., J.T.W, T.S.; Visualization: E.T., J.T.W.; Supervision: M.S.C.; Project administration: M.S.C.; Funding acquisition: M.S.C.

## Data availability

MS DDA Data has been deposited as a complete submission to the MassIVE repository (https://massive.ucsd.edu/ProteoSAFe/static/massive.jsp) and assigned the accession number MSV000089647. The ProteomeXchange accession is PXD034502. The dataset is currently available for reviewers at ftp://MSV000089647@massive.ucsd.edu. Please login with username MSV000089647reviewer; password: POC5.

MS DIA Data has been deposited as a complete submission to the MassIVE repository (https://massive.ucsd.edu/ProteoSAFe/static/massive.jsp) and assigned the accession number MSV000089651. The ProteomeXchange accession is PXD034506. The dataset is currently available for reviewers at ftp://MSV000089651@massive.ucsd.edu. Please login with username MSV000089651_reviewer; password: POC5. Both datasets will be made public upon acceptance of the manuscript.

---

**Fig. S1.**
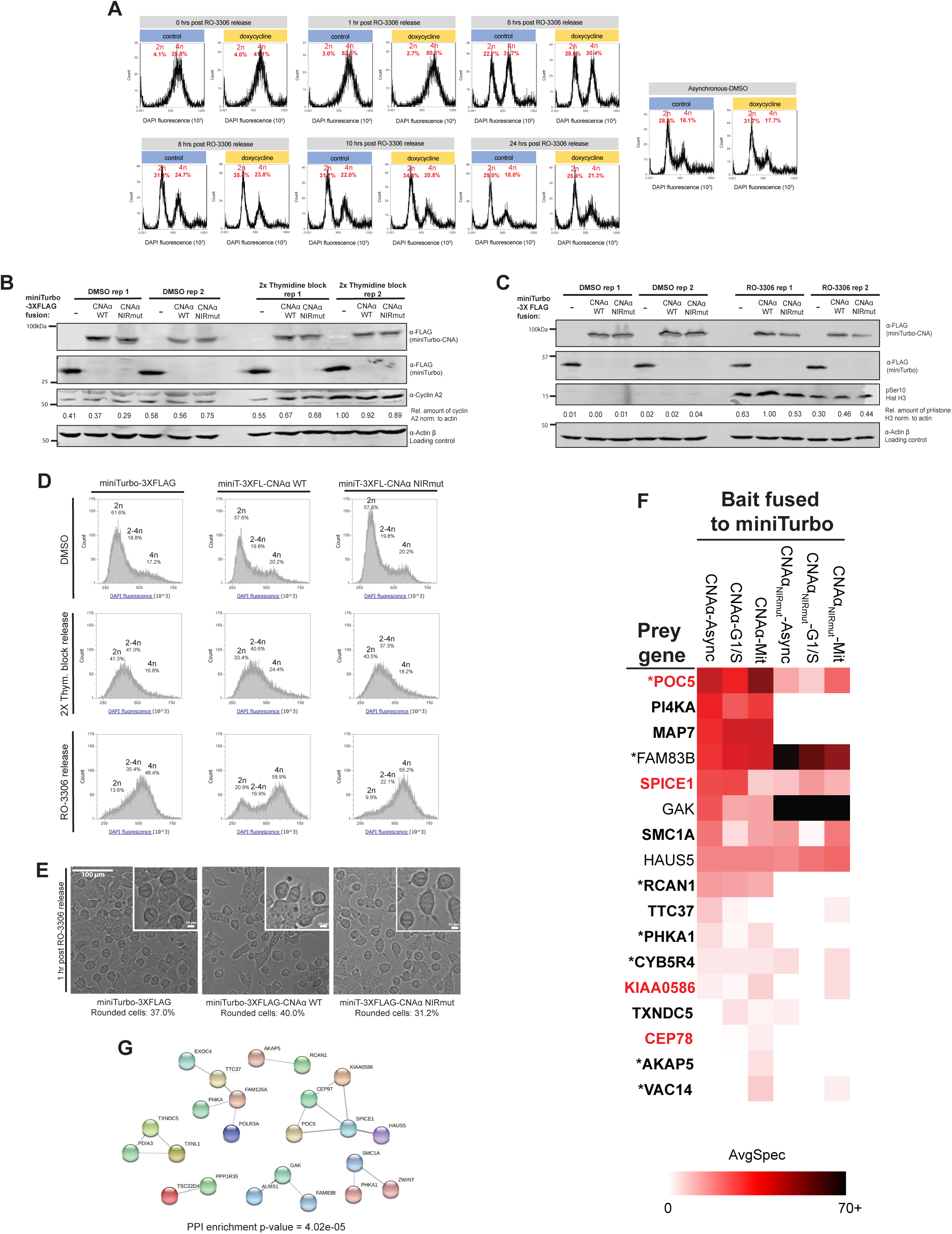
(Related to Figure 1). Proximity-labelled cells showed equal levels of miniTurbo bait expression, proper cell cycle synchronization and resulted in preys known to associate with one another. A. Induction of miniTurbo-CNAα overexpression does not disrupt cell cycle progression. HEK293 Flp-In T-Rex with inducible miniTurbo-CNAα_WT_ expression were either treated with 0 μg/mL doxycycline (control-blue boxes) or 1 μg/mL doxycycline (doxycycline-yellow boxes) for 48 hours. On the second day of doxycycline induction, cells were also treated with DMSO (asynchronous) or 9 μM RO-3306 for 20 hours and then released from arrest. At various time points post RO-3306 washout, cells were fixed, stained with 20 μg/mL DAPI and analyzed by flow cytometry. Cell cycle profiles post-release were similar between the control and doxycycline groups. B. Protein expression in asynchronous and G1/S miniTurbo samples. HEK293 Flp-In T-Rex cells with induced expression of miniTurbo fusions were synchronized and labelled with biotin. Immunoblot contains samples from two independent experimental replicates. Anti-FLAG staining was used to ensure expression of the miniTurbo baits, anti-cyclin A2 as a cell cycle marker and anti-actin beta as a loading control. Rep, replicate. Rel. amount of cyclin A2 norm. to actin: relative ratio of cyclin A2 band signal normalized by the corresponding actin band signal. C. Protein expression in asynchronous and mitotic miniTurbo samples. HEK293 Flp-In T-Rex cells with induced expression of miniTurbo fusions were synchronized and labelled with biotin. Immunoblot contains samples from two independent experimental replicates. Anti-FLAG staining was used to ensure expression of the miniTurbo baits, anti-Ser10 phosphorylated histone H3 as a mitotic marker and anti-actin beta as a loading control. Rep, replicate. Rel. amount of pHistone H3 norm. to actin: relative ratio of phosphorylated histone H3 band signal normalized by the corresponding actin band signal. D. Cell cycle profiles of synchronized miniTurbo HEK293 Flp-In T-Rex cells prior to mass spectrometry analysis. Profiles shown are from one of the two independent replicates of the proximity labeling experiment and were obtained by flow cytometry using DAPI staining. E. Brightfield microscopy images showing miniTurbo cell samples synchronized in mitosis 1 hour after RO-3306 washout. Insets show that cells are rounded up and their chromosomes are aligned on the metaphase plate, indicating that they are mitotic. Scale bar, 100 μm, inset scale bar, 10 μm. F. Heat map of average spectral counts for preys labelled by miniTurbo-CNA_WT_ or CNA_NIRmut_ in asynchronous, G1/S and mitotic cells. Only preys labelled by CNA_WT_ with unique peptides ≥ 2 and bayesian false discovery rate (BFDR) ≤ 0.01 in at least one treatment are included. Spectral counts were obtained via data-independent mixture-spectrum partitioning using libraries of identified tandem mass spectra (DIA/mSPLIT). Bold: preys with PxIxIT-dependent biotinylation (Log_2_ CNA_WT/_CNA_NIRmut_ ≥ 0.5). Red: proteins annotated with the Gene Ontology (GO) term “centrosome”. Asterisks: proteins with predicted CN-dependent SLiMs (PxIxIT or LxVP) identified *in silico* (Wigington et al., 2020). AvgSpec, average spectral counts. Exact spectral counts can be found under “mSPLIT filtered dataset” on Table S1. G. STRING database v11.5 (string-db.org) network of protein-protein interactions between CN-proximal proteins identified in this study (38 from DDA analysis and 3 additional from mSPLIT analysis). Proteins without any known interactors within the network were omitted. Line intensity is proportional to the confidence of the interaction. PPI enrichment p-value: 4.02e-05. Number of edges: 19. Expected number of edges: 6.

**Fig. S2.**
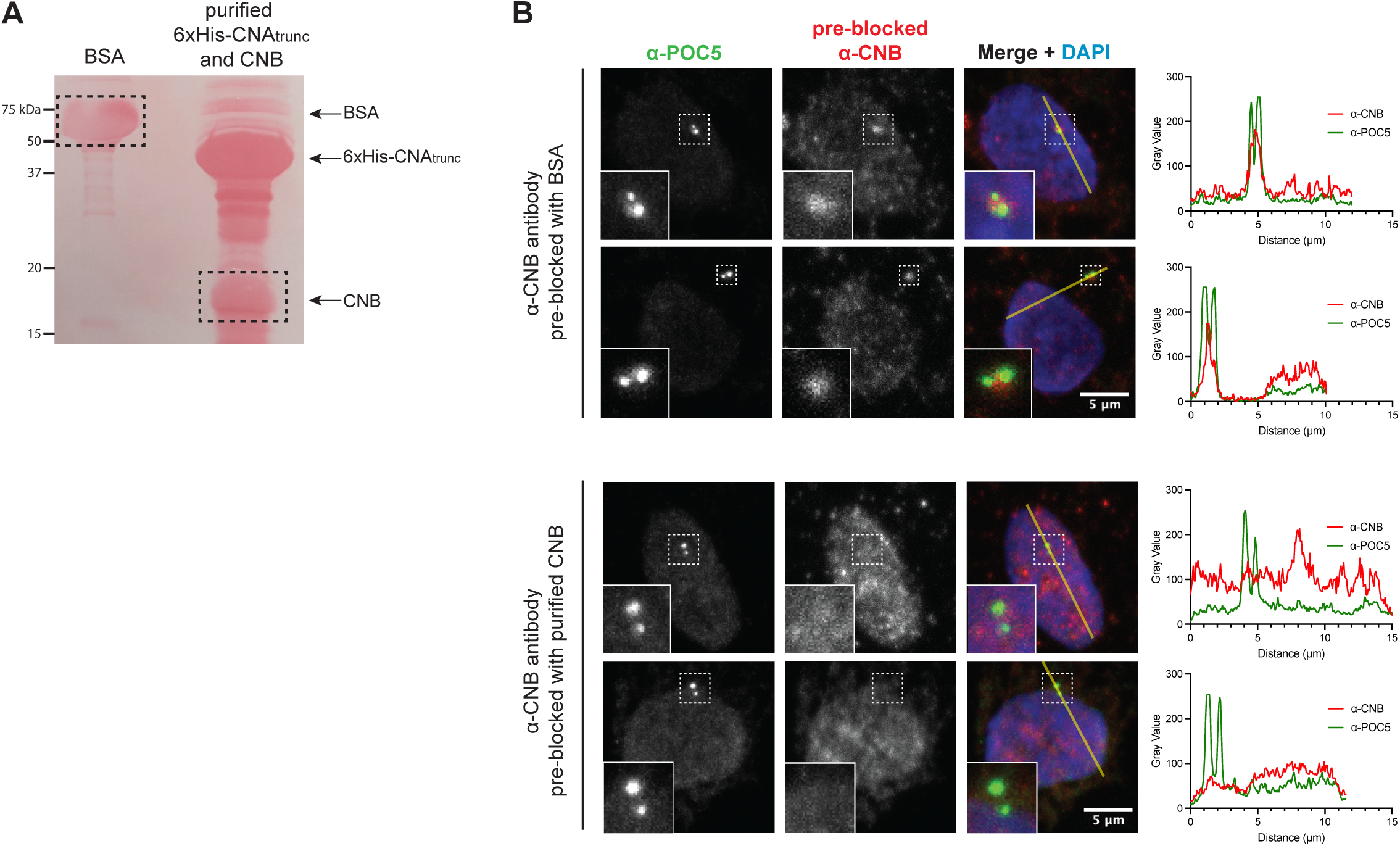
(Related to Figure 2) CNB antibody blocking demonstrates that centrosomal localization of CNB is highly specific. A. Nitrocellulose membrane stained with Ponceau S, showing analysis of purified BSA and purified truncated CNA and CNB by SDS-PAGE. Dotted lines indicate pieces of the membrane that were excised and incubated with anti-CNB antibody for antibody blocking in Fig. S2B. B. Pre-incubating anti-CNB antibody with purified CNB eliminates centrosomal localization. Immunofluorescence of cytosol depleted hTERT-RPE1 cells. Centrioles are marked by anti-POC5 staining (green) and nuclei are marked by DAPI (blue). CNB localization (red) is analyzed by staining cells with anti-CNB antibody that has been pre-incubated with purified proteins transferred onto a nitrocellulose membrane as shown in Fig. S2A. In the top panels, anti-CNB antibody was incubated with bovine serum albumin (BSA) in two independent experiments. In the bottom panels, anti-CNB antibody was incubated with CNB in two independent experiments. Lines were drawn across the two centrioles of each cell (shown in yellow) to generate line intensity plots on the right of each immunofluorescence panel. Line intensity plots track the intensity of CNB signal (red) and POC5 signal (green) across the cell and the two centrioles, indicated by the double peaks of POC5 intensity. Scale bar, 5 μm.

**Fig. S3.**
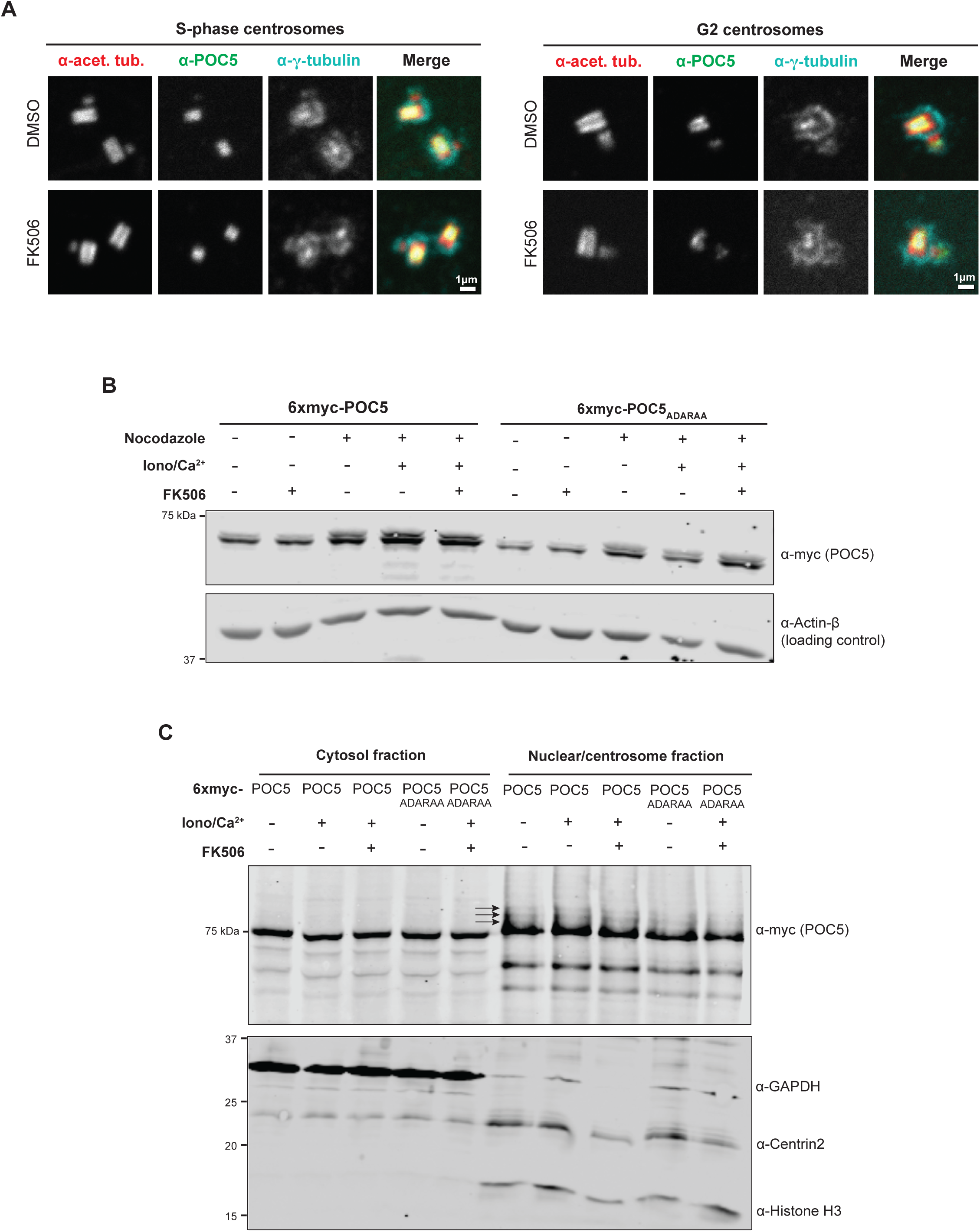
(Related to Figure 3) Calcineurin activity disrupts POC5 distribution, but does not alter γ-tubulin distribution or POC5 phospho-status *in vivo*. A. Expansion microscopy of S-phase and G2 hTERT-RPE1 cells treated with DMSO or FK506 for 48 hours, showing POC5 in the centriole lumen and γ-tubulin in the lumen and PCM. Images obtained from a single z-plane. Scale bar, 1 μm. B. POC5 is phosphorylated in mitosis independently of CN stimulation or inhibition. Immunoblot showing lysates of HeLa cells transfected with 6xmyc-POC5 or -POC5_ADARAA._ Cells were treated with DMSO (-) or 100 ng nocodazole (+) for 18 hours and then incubated for one additional hour in 37°C after drug washout. The second POC5 band that appears in nocodazole (+) samples corresponds to p-POC5. For CN activation, samples were additionally treated with 1 μM ionomycin + 1 mM CaCl_2_ for one hour prior to cell lysis. For CN inhibition, samples were treated with 2.5 μM FK506 for one hour followed by 2.5 μM FK506 + 1 μM ionomycin + 1 mM CaCl_2_ for one more hour prior to cell lysis. Iono, ionomycin. Ca^2+^, calcium ions from CaCl_2_ addition. Anti-actin-beta was used as a loading control. C. POC5 is hyperphosphorylated at centrosomes independently of CN stimulation or inhibition. Cytosolic and nuclear-centrosomal fractions prepared via sucrose fractionation from HeLa cells transfected with 6xmyc-POC5 or -POC5_ADARAA_. For CN activation, samples were treated with 1 μM ionomycin + 1 mM CaCl_2_ for one hour prior to fractionation. For CN inhibition, samples were treated with 2.5 μM FK506 for one hour followed by 2.5 μM FK506 + 1 μM ionomycin + 1 mM CaCl_2_ for one more hour prior to fractionation. Iono, ionomycin. Ca^2+^, calcium ions from CaCl_2_ addition. Anti-GAPDH was used a cytosolic marker, anti-centrin2 as a centrosomal marker and anti-histone H3 as a nuclear marker.

**Fig. S4.**
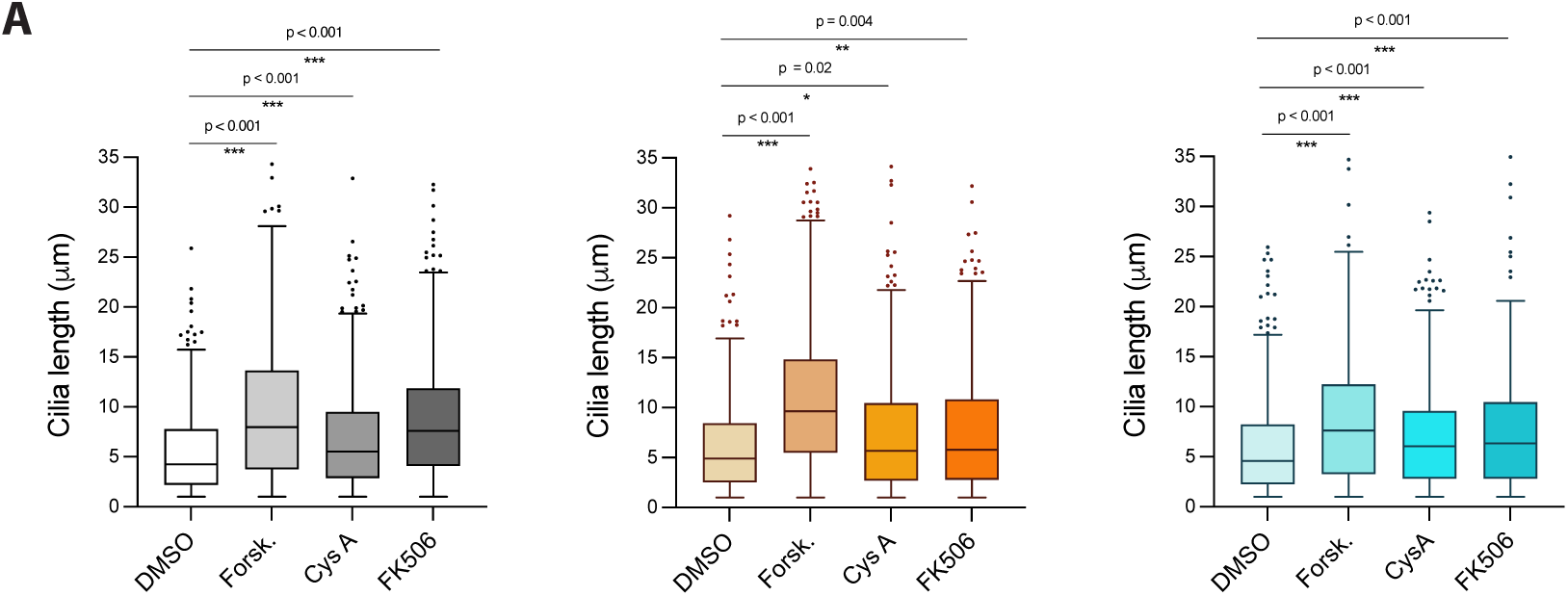
(Related to Figure 4) Calcineurin inhibition increases cilia length. A. Forskolin treatment and CN inhibition consistently promote cilia elongation. Cilia length in ciliated IMCD3 cells treated for 3 hours with DMSO, forskolin, cyclosporin A or FK506 as in Fig. 4A. Graphs show data from three independent experimental replicates in addition to the replicate shown in 4B. Cilia length was determined by 3D vector analysis of confocal z-stacks using CiliaQ with CANNY 3D segmentation (Hansen et al., 2021). Only continuous Arl13b branches with length > 1 μm are shown on the graph. Number of cilia measured: leftmost graph, DMSO, n=475, forskolin, n=564, cyclosporin A, n=481, FK506, n=718. Center graph, DMSO, n=497, forskolin, n=602, cyclosporin A, n=603, FK506, n=494. Rightmost graph, DMSO, n=371, forskolin, n=417, cyclosporin A, n=421, FK506, n=470. Boxplots show median length ± interquartile range (IQR) and whiskers represent the median ± 1.5 x IQR. n.s., not significant, *p<0.05, **p<0.01, ***p<0.001. P-values are indicated on each graph, using two-tailed Mann-Whitney test.

**Table S1. Dataset of proteins identified via PDB-MS, final filtered dataset and associated GO term analyses.**

MiniTurbo-MS data-dependent acquisition (DDA) dataset analyzed by SAINTexp3.6.1 and data-independent mixture-spectrum partitioning using libraries of identified tandem mass spectra (mSPLIT) dataset analyzed by SAINTexp3.6.3, filtered DDA dataset and associated GO Term analyses.

**Table S2.** Subcellular localization of centrosome-associated, calcineurin-proximal proteins. Table of relevant literature regarding cilia and centrosome components that were proximal to miniTurbo-CN in our study. Proteins are identified by their corresponding gene name, with a brief summary of their precise subcellular location as described by the referenced studies.

| Gene name | Protein localization | References |
| --- | --- | --- |
| AKAP5 (AKAP79, AKAP150) | Cilia, centrosomes, spindle poles | (Choi et al., 2011) |
| ALMS1 | Proximal centrioles | (Knorz et al., 2010) |
| CDC16 | Centrosomes and spindle poles | (Tugendreich et al., 1995) |
| CEP78 | Distal centrioles | (Hossain et al., 2017) |
| CEP97 | Distal centrioles | (Spektor et al., 2007) |
| EXOC4 (SEC8) | Cilia, centrosomes | (Gupta et al., 2015; Zuo et al., 2011) |
| GAK | Spindle poles during mitosis | (Fukushima et al., 2017; Naito et al., 2012) |
| HAUS5 | Centrosomes in interphase, mitotic spindle microtubules, middle to distal centrioles | (Lawo et al., 2009; Schweizer et al., 2021) |
| HOOK1 | Broad distribution around centrosomes | (Szebenyi et al., 2007) |
| POC5 | Middle to distal centrioles | (Le Guennec et al., 2020) |
| PPP1R35 | Proximal centrioles | (Sydor et al., 2018) |
| SMC1A | Centrioles, spindle poles | (Guan et al., 2008; Wong, 2010) |
| SPICE1 | Proximal centrioles (distal to SAS-6, proximal to centrin) | (Comartin et al., 2013) |
| TALPID3 (KIAA0586) | Distal centrioles | (Kobayashi et al., 2014) |

## References

1. Almeida, J. L., Dakic, A., Kindig, K., Kone, M., Letham, D. L. D., Langdon, S., Peat, R., Holding-Pillai, J., Hall, E. M., Ladd, M., et al. (2019). Interlaboratory study to validate a STR profiling method for intraspecies identification of mouse cell lines. PLoS One 14, e0218412.

2. Avasthi, P. and Marshall, W. F. (2012). Stages of ciliogenesis and regulation of ciliary length. Differentiation 83, S30–42.

3. Azimzadeh, J., Hergert, P., Delouvée, A., Euteneuer, U., Formstecher, E., Khodjakov, A. and Bornens, M. (2009). hPOC5 is a centrin-binding protein required for assembly of full-length centrioles. J. Cell Biol. 185, 101–114.

4. Azzi, J. R., Sayegh, M. H. and Mallat, S. G. (2013). Calcineurin inhibitors: 40 years later, can’t live without. J. Immunol. 191, 5785–5791.

5. Barallon, R., Bauer, S. R., Butler, J., Capes-Davis, A., Dirks, W. G., Elmore, E., Furtado, M., Kline, M. C., Kohara, A., Los, G. V., et al. (2010). Recommendation of short tandem repeat profiling for authenticating human cell lines, stem cells, and tissues. In Vitro Cell. Dev. Biol. Anim. 46, 727–732.

6. Berridge, M. J., Bootman, M. D. and Roderick, H. L. (2003). Calcium signalling: dynamics, homeostasis and remodelling. Nat. Rev. Mol. Cell Biol. 4, 517–529.

7. Besschetnova, T. Y., Kolpakova-Hart, E., Guan, Y., Zhou, J., Olsen, B. R. and Shah, J. V. (2010). Identification of signaling pathways regulating primary cilium length and flow-mediated adaptation. Curr. Biol. 20, 182–187.

8. Branon, T. C., Bosch, J. A., Sanchez, A. D., Udeshi, N. D., Svinkina, T., Carr, S. A., Feldman, J. L., Perrimon, N. and Ting, A. Y. (2018). Efficient proximity labeling in living cells and organisms with TurboID. Nat. Biotechnol. 36, 880–887.

9. Brauer, B. L., Moon, T. M., Sheftic, S. R., Nasa, I., Page, R., Peti, W. and Kettenbach, A. N. (2019). Leveraging new definitions of the LxVP SLiM to discover novel calcineurin regulators and substrates. ACS Chem. Biol. 14, 2672– 2682.

10. Choi, Y.-H., Suzuki, A., Hajarnis, S., Ma, Z., Chapin, H. C., Caplan, M. J., Pontoglio, M., Somlo, S. and Igarashi, P. (2011). Polycystin-2 and phosphodiesterase 4C are components of a ciliary A-kinase anchoring protein complex that is disrupted in cystic kidney diseases. Proc. Natl. Acad. Sci. U. S. A. 108, 10679–10684.

11. Comartin, D., Gupta, G. D., Fussner, E., Coyaud, É., Hasegan, M., Archinti, M., Cheung, S. W. T., Pinchev, D., Lawo, S., Raught, B., et al. (2013). CEP120 and SPICE1 cooperate with CPAP in centriole elongation. Curr. Biol. 23, 1360– 1366.

12. Dell’Acqua, M. L., Dodge, K. L., Tavalin, S. J. and Scott, J. D. (2002). Mapping the protein phosphatase-2B anchoring site on AKAP79. J. Biol. Chem. 277, 48796– 48802.

13. Delling, M., DeCaen, P. G., Doerner, J. F., Febvay, S. and Clapham, D. E. (2013). Primary cilia are specialized calcium signalling organelles. Nature 504, 311–314.

14. Eng, J. K., Jahan, T. A. and Hoopmann, M. R. (2013). Comet: an open-source MS/MS sequence database search tool. Proteomics 13, 22–24.

15. Farouk, S. S. and Rein, J. L. (2020). The many faces of calcineurin inhibitor toxicity-what the FK? Adv. Chronic Kidney Dis. 27, 56–66.

16. Fukushima, K., Wang, M., Naito, Y., Uchihashi, T., Kato, Y., Mukai, S., Yabuta, N. and Nojima, H. (2017). GAK is phosphorylated by c-Src and translocated from the centrosome to chromatin at the end of telophase. Cell Cycle 16, 415–427.

17. Gambarotto, D., Zwettler, F. U., Le Guennec, M., Schmidt-Cernohorska, M., Fortun, D., Borgers, S., Heine, J., Schloetel, J.-G., Reuss, M., Unser, M., et al. (2019). Imaging cellular ultrastructures using expansion microscopy (U-ExM). Nat. Methods 16, 71–74.

18. Gingras, A.-C., Abe, K. T. and Raught, B. (2019). Getting to know the neighborhood: using proximity-dependent biotinylation to characterize protein complexes and map organelles. Curr. Opin. Chem. Biol. 48, 44–54.

19. Grigoriu, S., Bond, R., Cossio, P., Chen, J. A., Ly, N., Hummer, G., Page, R., Cyert, M. S. and Peti, W. (2013). The molecular mechanism of substrate engagement and immunosuppressant inhibition of calcineurin. PLoS Biol. 11, e1001492.

20. Guan, J., Ekwurtzel, E., Kvist, U. and Yuan, L. (2008). Cohesin protein SMC1 is a centrosomal protein. Biochem. Biophys. Res. Commun. 372, 761–764.

21. Gupta, G. D., Coyaud, É., Gonçalves, J., Mojarad, B. A., Liu, Y., Wu, Q., Gheiratmand, L., Comartin, D., Tkach, J. M., Cheung, S. W. T., et al. (2015). A dynamic protein interaction landscape of the human centrosome-cilium interface. Cell 163, 1484–1499.

22. Hansen, J. N., Rassmann, S., Stüven, B., Jurisch-Yaksi, N. and Wachten, D. (2021). CiliaQ: a simple, open-source software for automated quantification of ciliary morphology and fluorescence in 2D, 3D, and 4D images. Eur. Phys. J. E Soft Matter 44, 18.

23. Hassan, A., Parent, S., Mathieu, H., Zaouter, C., Molidperee, S., Bagu, E. T., Barchi, S., Villemure, I., Patten, S. A. and Moldovan, F. (2019). Adolescent idiopathic scoliosis associated POC5 mutation impairs cell cycle, cilia length and centrosome protein interactions. PLoS One 14, e0213269.

24. Helassa, N., Nugues, C., Rajamanoharan, D., Burgoyne, R. D. and Haynes, L. P. (2019). A centrosome-localized calcium signal is essential for mammalian cell mitosis. FASEB J. 33, 14602–14610.

25. Hesketh, G. G., Youn, J.-Y., Samavarchi-Tehrani, P., Raught, B. and Gingras, A.-C. (2017). Parallel exploration of interaction space by BioID and affinity purification coupled to mass spectrometry. Methods Mol. Biol. 1550, 115–136.

26. Hildebrandt, F., Benzing, T. and Katsanis, N. (2011). Ciliopathies. N. Engl. J. Med. 364, 1533–1543.

27. Hornbeck, P. V., Zhang, B., Murray, B., Kornhauser, J. M., Latham, V. and Skrzypek, E. (2015). PhosphoSitePlus, 2014: mutations, PTMs and recalibrations. Nucleic Acids Res. 43, D512–20.

28. Hossain, D., Javadi Esfehani, Y., Das, A. and Tsang, W. Y. (2017). Cep78 controls centrosome homeostasis by inhibiting EDD-DYRK2-DDB1VprBP. EMBO Rep. 18, 632–644.

29. Hu, J., Bae, Y.-K., Knobel, K. M. and Barr, M. M. (2006). Casein kinase II and calcineurin modulate TRPP function and ciliary localization. Mol. Biol. Cell 17, 2200–2211.

30. Ingebritsen, T. S. and Cohen, P. (1983). The protein phosphatases involved in cellular regulation. 1. Classification and substrate specificities. Eur. J. Biochem. 132, 255–261.

31. Joly, D., Ishibe, S., Nickel, C., Yu, Z., Somlo, S. and Cantley, L. G. (2006). The polycystin 1-C-terminal fragment stimulates ERK-dependent spreading of renal epithelial cells. J. Biol. Chem. 281, 26329–26339.

32. Kee, H. L., Dishinger, J. F., Blasius, T. L., Liu, C.-J., Margolis, B. and Verhey, K. J. (2012). A size-exclusion permeability barrier and nucleoporins characterize a ciliary pore complex that regulates transport into cilia. Nat. Cell Biol. 14, 431– 437.

33. Khouj, E. M., Prosser, S. L., Tada, H., Chong, W. M., Liao, J.-C., Sugasawa, K. and Morrison, C. G. (2019). Differential requirements for the EF-hand domains of human centrin 2 in primary ciliogenesis and nucleotide excision repair. J. Cell Sci. 132, jcs228486.

34. Knight, J. D. R., Liu, G., Zhang, J. P., Pasculescu, A., Choi, H. and Gingras, A.-C. (2015). A web-tool for visualizing quantitative protein-protein interaction data. Proteomics 15, 1432–1436.

35. Knorz, V. J., Spalluto, C., Lessard, M., Purvis, T. L., Adigun, F. F., Collin, G. B., Hanley, N. A., Wilson, D. I. and Hearn, T. (2010). Centriolar association of ALMS1 and likely centrosomal functions of the ALMS motif–containing proteins C10orf90 and KIAA1731. Mol. Biol. Cell 21, 3617–3629.

36. Kobayashi, T., Kim, S., Lin, Y.-C., Inoue, T. and Dynlacht, B. D. (2014). The CP110-interacting proteins Talpid3 and Cep290 play overlapping and distinct roles in cilia assembly. J. Cell Biol. 204, 215–229.

37. Lawo, S., Bashkurov, M., Mullin, M., Ferreria, M. G., Kittler, R., Habermann, B., Tagliaferro, A., Poser, I., Hutchins, J. R. A., Hegemann, B., et al. (2009). HAUS, the 8-subunit human Augmin complex, regulates centrosome and spindle integrity. Curr. Biol. 19, 816–826.

38. Le Guennec, M., Klena, N., Gambarotto, D., Laporte, M. H., Tassin, A.-M., van den Hoek, H., Erdmann, P. S., Schaffer, M., Kovacik, L., Borgers, S., et al. (2020). A helical inner scaffold provides a structural basis for centriole cohesion. Sci. Adv. 6, eaaz4137.

39. Li, H., Zhang, L., Rao, A., Harrison, S. C. and Hogan, P. G. (2007). Structure of calcineurin in complex with PVIVIT peptide: portrait of a low-affinity signalling interaction. J. Mol. Biol. 369, 1296–1306.

40. Li, S.-J., Wang, J., Ma, L., Lu, C., Wang, J., Wu, J.-W. and Wang, Z.-X. (2016). Cooperative autoinhibition and multi-level activation mechanisms of calcineurin. Cell Res. 26, 336–349.

41. Liu, G., Knight, J. D. R., Zhang, J. P., Tsou, C.-C., Wang, J., Lambert, J.-P., Larsen, B., Tyers, M., Raught, B., Bandeira, N., et al. (2016). Data Independent Acquisition analysis in ProHits 4.0. J. Proteomics 149, 64–68.

42. Mehta, S., Li, H., Hogan, P. G. and Cunningham, K. W. (2009). Domain architecture of the regulators of calcineurin (RCANs) and identification of a divergent RCAN in yeast. Mol. Cell. Biol. 29, 2777–2793.

43. Mi, H., Muruganujan, A., Casagrande, J. T. and Thomas, P. D. (2013). Large-scale gene function analysis with the PANTHER classification system. Nat. Protoc. 8, 1551–1566.

44. Moreno-Leon, L., West, E. L., O’Hara-Wright, M., Li, L., Nair, R., He, J., Anand, M., Sahu, B., Chavali, V. R. M., Smith, A. J., et al. (2021). RPGR isoform imbalance causes ciliary defects due to exon ORF15 mutations in X-linked retinitis pigmentosa (XLRP). Hum. Mol. Genet. 29, 3706–3716.

45. Naito, Y., Shimizu, H., Kasama, T., Sato, J., Tabara, H., Okamoto, A., Yabuta, N. and Nojima, H. (2012). Cyclin G-associated kinase regulates protein phosphatase 2A by phosphorylation of its B’γ subunit. Cell Cycle 11, 604–616.

46. Perkins, D. N., Pappin, D. J., Creasy, D. M. and Cottrell, J. S. (1999). Probability-based protein identification by searching sequence databases using mass spectrometry data. Electrophoresis 20, 3551–3567.

47. Plotnikova, O. V., Nikonova, A. S., Loskutov, Y. V., Kozyulina, P. Y., Pugacheva, E. N. and Golemis, E. A. (2012). Calmodulin activation of Aurora-A kinase (AURKA) is required during ciliary disassembly and in mitosis. Mol. Biol. Cell 23, 2658–2670.

48. Roy, J. and Cyert, M. S. (2020). Identifying new substrates and functions for an old enzyme: Calcineurin. Cold Spring Harb. Perspect. Biol. 12, a035436.

49. Rusnak, F. and Mertz, P. (2000). Calcineurin: Form and function. Physiol. Rev. 80, 1483–1521.

50. Schweizer, N., Haren, L., Dutto, I., Viais, R., Lacasa, C., Merdes, A. and Lüders, J. (2021). Sub-centrosomal mapping identifies augmin-γTuRC as part of a centriole-stabilizing scaffold. Nat. Commun. 12, 6042.

51. Sheftic, S. R., Page, R. and Peti, W. (2016). Investigating the human Calcineurin Interaction Network using the πɸLxVP SLiM. Sci. Rep. 6, 38920.

52. Shteynberg, D., Deutsch, E. W., Lam, H., Eng, J. K., Sun, Z., Tasman, N., Mendoza, L., Moritz, R. L., Aebersold, R. and Nesvizhskii, A. I. (2011). iProphet: multi-level integrative analysis of shotgun proteomic data improves peptide and protein identification rates and error estimates. Mol. Cell. Proteomics 10, M111.007690.

53. Spektor, A., Tsang, W. Y., Khoo, D. and Dynlacht, B. D. (2007). Cep97 and CP110 suppress a cilia assembly program. Cell 130, 678–690.

54. Stevenson, N. L., Bergen, D. J. M., Xu, A., Wyatt, E., Henry, F., McCaughey, J., Vuolo, L., Hammond, C. L. and Stephens, D. J. (2018). Regulator of calcineurin-2 is a centriolar protein with a role in cilia length control. J. Cell Sci. 131, jcs212258.

55. Sydor, A. M., Coyaud, E., Rovelli, C., Laurent, E., Liu, H., Raught, B. and Mennella, V. (2018). PPP1R35 is a novel centrosomal protein that regulates centriole length in concert with the microcephaly protein RTTN. Elife 7,.

56. Szebenyi, G., Hall, B., Yu, R., Hashim, A. I. and Krämer, H. (2007). Hook2 localizes to the centrosome, binds directly to centriolin/CEP110 and contributes to centrosomal function. Traffic 8, 32–46.

57. Teo, G., Liu, G., Zhang, J., Nesvizhskii, A. I., Gingras, A.-C. and Choi, H. (2014). SAINTexpress: improvements and additional features in Significance Analysis of INTeractome software. J. Proteomics 100, 37–43.

58. Tompa, P., Davey, N. E., Gibson, T. J. and Babu, M. M. (2014). A million peptide motifs for the molecular biologist. Mol. Cell 55, 161–169.

59. Tugendreich, S., Tomkiel, J., Earnshaw, W. and Hieter, P. (1995). CDC27Hs colocalizes with CDC16Hs to the centrosome and mitotic spindle and is essential for the metaphase to anaphase transition. Cell 81, 261–268.

60. Ulengin-Talkish, I., Parson, M. A. H., Jenkins, M. L., Roy, J., Shih, A. Z. L., St-Denis, N., Gulyas, G., Balla, T., Gingras, A.-C., Várnai, P., et al. (2021). Palmitoylation targets the calcineurin phosphatase to the phosphatidylinositol 4-kinase complex at the plasma membrane. Nat. Commun. 12, 6064.

61. Veland, I. R., Awan, A., Pedersen, L. B., Yoder, B. K. and Christensen, S. T. (2009). Primary cilia and signaling pathways in mammalian development, health and disease. Nephron Physiol. 111, 39–53.

62. Wang, J. T. and Stearns, T. (2017). The ABCs of centriole architecture: The form and function of triplet microtubules. Cold Spring Harb. Symp. Quant. Biol. 82, 145– 155.

63. Wang, J., Tucholska, M., Knight, J. D. R., Lambert, J.-P., Tate, S., Larsen, B., Gingras, A.-C. and Bandeira, N. (2015). MSPLIT-DIA: sensitive peptide identification for data-independent acquisition. Nat. Methods 12, 1106–1108.

64. Weisz Hubshman, M., Broekman, S., van Wijk, E., Cremers, F., Abu-Diab, A., Khateb, S., Tzur, S., Lagovsky, I., Smirin-Yosef, P., Sharon, D., et al. (2018). Whole-exome sequencing reveals POC5 as a novel gene associated with autosomal recessive retinitis pigmentosa. Hum. Mol. Genet. 27, 614–624.

65. Wigington, C. P., Roy, J., Damle, N. P., Yadav, V. K., Blikstad, C., Resch, E., Wong, C. J., Mackay, D. R., Wang, J. T., Krystkowiak, I., et al. (2020). Systematic discovery of short linear motifs decodes calcineurin phosphatase signaling. Mol. Cell 79, 342–358.e12.

66. Wong, R. W. (2010). Interaction between Rae1 and cohesin subunit SMC1 is required for proper spindle formation. Cell Cycle 9, 198–200.

67. Zuo, X., Fogelgren, B. and Lipschutz, J. H. (2011). The small GTPase Cdc42 is necessary for primary ciliogenesis in renal tubular epithelial cells. J. Biol. Chem. 286, 22469–22477.

